# Physical mechanisms of red blood cell splenic filtration

**DOI:** 10.1101/2023.01.10.523245

**Authors:** Alexis Moreau, François Yaya, Huijie Lu, Anagha Surendranath, Anne Charrier, Benoit Dehapiot, Emmanuèle Helfer, Annie Viallat, Zhangli Peng

## Abstract

The splenic interendothelial slits fulfill the essential function of continuously filtering red blood cells (RBCs) from the bloodstream to eliminate abnormal and aged cells. To date, the process by which 8 µm RBCs pass through 0.3 µm-wide slits remains enigmatic. Does the slit caliber increase during RBC passage as sometimes suggested? Here, we elucidated the mechanisms that govern the RBC retention or passage dynamics in slits by combining multiscale modeling, live imaging, and microfluidic experiments on an original device with sub-micron wide physiologically calibrated slits. We observed that healthy RBCs pass through 0.28 µm-wide rigid slits at 37°C. To achieve this feat, they must meet two requirements. Geometrically, their surface area-to-volume ratio must be compatible with a shape in two tether-connected equal spheres. Mechanically, the cells with a low surface area-to-volume ratio (28 % of RBCs in a 0.4 µm-wide slit) must locally unfold their spectrin cytoskeleton inside the slit. In contrast, activation of the mechanosensitive PIEZO1 channel is not required. The RBC transit time through the slits follows a -1 and -3 power law with in-slit pressure drop and slip width, respectively. This law is similar to that of a Newtonian fluid in a 2D Poiseuille flow, showing that the dynamics of RBCs is controlled by their cytoplasmic viscosity. Altogether, our results show that filtration through submicron-wide slits is possible without further slit opening. Furthermore, our approach addresses the critical need for in-vitro evaluation of splenic clearance of diseased or engineered RBCs for transfusion and drug delivery.

**Significance Statement:** Splenic filtration of red blood cells through narrow interendothelial slits remains poorly understood despite its physiological significance as experiments and imaging of red cells passing through the slits are lacking. Here, we coupled live imaging, biomimetic submicron-fluidics, and multiscale modeling to quantify passage conditions. Remarkably, healthy 8-µm cells can pass through 0.28-µm slits at body temperature. This event is conditioned to cells being able to deform into two tether-connected equal spheres and, in limiting cases, to unfold their spectrin cytoskeleton. We showed that cells behave like a Newtonian fluid and that their dynamics is controlled by the inner fluid viscosity. We thus propose an in-vitro and in-silico approach to quantify splenic clearance of diseased cells and cells engineered for transfusion and drug delivery.

---

In the microcirculation, red blood cells (RBCs) must deform strongly to squeeze into microcapillaries smaller than themselves. This requires very specific morphological and mechanical cellular properties. Indeed, an RBC is a biconcave disk-shaped capsule with a large surface area-to-volume ratio. Its shell is a membrane composed of an outer fluid inextensible lipid bilayer and an inner cytoskeleton made of spectrin tetramers forming a two-dimensional viscoelastic network (1, 2). Its inner volume is a solution of hemoglobin. During the four months of RBC life, both their surface area-to-volume ratio and cytoskeleton’s viscoelasticity evolve resulting in a progressive degradation of the cell deformability (3). Moreover, in many diseases, such as sickle cell disease or malaria infection, RBCs are less deformable (4, 5) and may have difficulty in squeezing in the narrowest blood capillaries. It is therefore essential that our body regularly tests the deformability of RBCs in order to eliminate damaged ones. The RBC selection occurs in the sinusoids of the red pulp of the spleen that each RBC visits approximatively every 200 minutes (6, 7). To return to the vascular system, RBCs must squeeze through slits of 0.25-1.2 µm in width, 0.9 -3.2 µm in length, and around 5 µm in depth (8, 9) between neighboring cells of the vein endothelium where they undergo extreme deformation as shown in electron microscopy images of RBCs in splenic slits (9, 10). This stringent RBC fitness test is fundamental to retain and eliminate unfit cells, and thus ensure proper blood microcirculation (8, 9, 11, 12). However, the values of the mechanical quantities selected during the passage of RBCs in the splenic slits and the associated dynamics, such as deformation mechanisms and transit times, remain poorly known today. Indeed, on the one hand, comprehensive in-vivo studies cannot be conducted because they are too invasive, and on the other hand, in-vitro studies are limited because the usual manufacturing methods do not allow the production of slits less than 2 µm in width, much larger than the physiological values (13–15). Stacks of beads may provide slits of less than 1 µm in width but do not allow controlled and extensive studies (11). Recent numerical and/or theoretical approaches have suggested that the spleen may play an important role in defining the surface area-to-volume ratio of the RBCs circulating in the microvascular system (16–19), but, so far, numerical approaches are not quantitatively validated by experiments and no experimental direct observation of RBCs flowing in splenic-like slits supports this hypothesis. Moreover, observations (20, 21) also suggest that slit caliber may vary over time leading to the hypothesis and modeling (22) that it is modulated by stress fiber contraction in human endothelial cells.

Here, we couple a unique in-vitro microfluidic technique to a multiscale in-silico RBC model that enables a quantitative approach of the mechanisms of passage of RBCs through interendothelial slits. The in-vitro technique allows the observation of the dynamics of passage of RBCs in slits of physiological dimensions, namely a submicron width, under tunable external stresses and slit sizes. The in-silico RBC model is implemented in a dynamic and a quasi-static versions. The dynamic version is integrated with a boundary integral simulation of surrounding flows to resolve the full fluid-cell interactions during this passage process, while the quasi-static version is done in commercial software ABAQUS (23) (see Materials and Methods and *SI Appendix*, Section 2).

We showed that red blood cells are capable of amazing extreme deformations allowing them to pass through rigid slits as narrow as 0.28 µm under a pressure drop of 500 Pa at body temperature, but not at room temperature. We first showed that, to cross the slits, the surface/volume ratio of individual cells must be sufficient to form two tether-connected equal spheres. Second, there is a threshold in the surface area-to-volume ratio, set by slit dimensions, in-slit pressure drop and temperature, below which RBCs must locally remodel by unfolding their spectrin network to pass through the slit. Finally, we showed that the transit time is mainly governed by the cytoplasmic viscosity and we quantitatively predicted this time as a function of the cell mechanical properties and external parameters: pressure drop, slit size, and temperature.

## Results

### In-vitro and in-silico observations: a quantitative approach

The in-vitro experiments and in-silico simulations were performed by varying the size of the slits and the pressure drop applied across the slits in the physiological range (9, 24). In-vivo measurements by Atkinson et al. (24) showed that intrasinusoidal pressures ranged between 260 and 1600 Pa. Accordingly, the in-slit pressure drop was varied between 200 and 2000 Pa and the width *W*, length *L* and depth *D* of the slits were chosen in the range 0.28-1.1 µm, 1.3-3 µm, 4.7-5.4 µm, respectively. The temperature was varied between 15°C and 37°C. All performed experiments are summarized in *SI Appendix*, Table S1. Experiments are illustrated in Fig. 1 and in *SI Appendix*, Movies S1 to S6. Figure 1A shows the schematics of an RBC approaching a slit entrance. Figure 1B shows first an electronic microscopy image of a 0.28-µm wide bridge on which a 0.28-µm wide slit is molded, and superimposed images extracted from *SI Appendix*, Movie S1 that illustrate the typical behavior of RBCs in slits. After exit, RBCs relax to their initial shape without apparent damage. Figure 1C discloses experimental timelapses showing two transiting cells, one with a classical dumbbell shape within the slit and the second one presenting a transient tip at its front, together with numerical simulations (see *SI Appendix*, Movies S2 and S3 of typical RBCs with dumbbell and tip shapes, respectively). The agreement between experiments and numerical simulations is remarkable, both on the similarity of cell shapes and on the full transit time of the cells. For each experiment several tens of RBCs were observed in the slits, allowing to measure each cell shape and transit time, and to quantify the retention rate of the observed RBC population. Most numerical simulations were done using an RBC standard shape, which is obtained by analyzing all the volumes and surface areas measured by many groups in the literature (*SI Appendix*, Table S2).

**Fig. 1.**
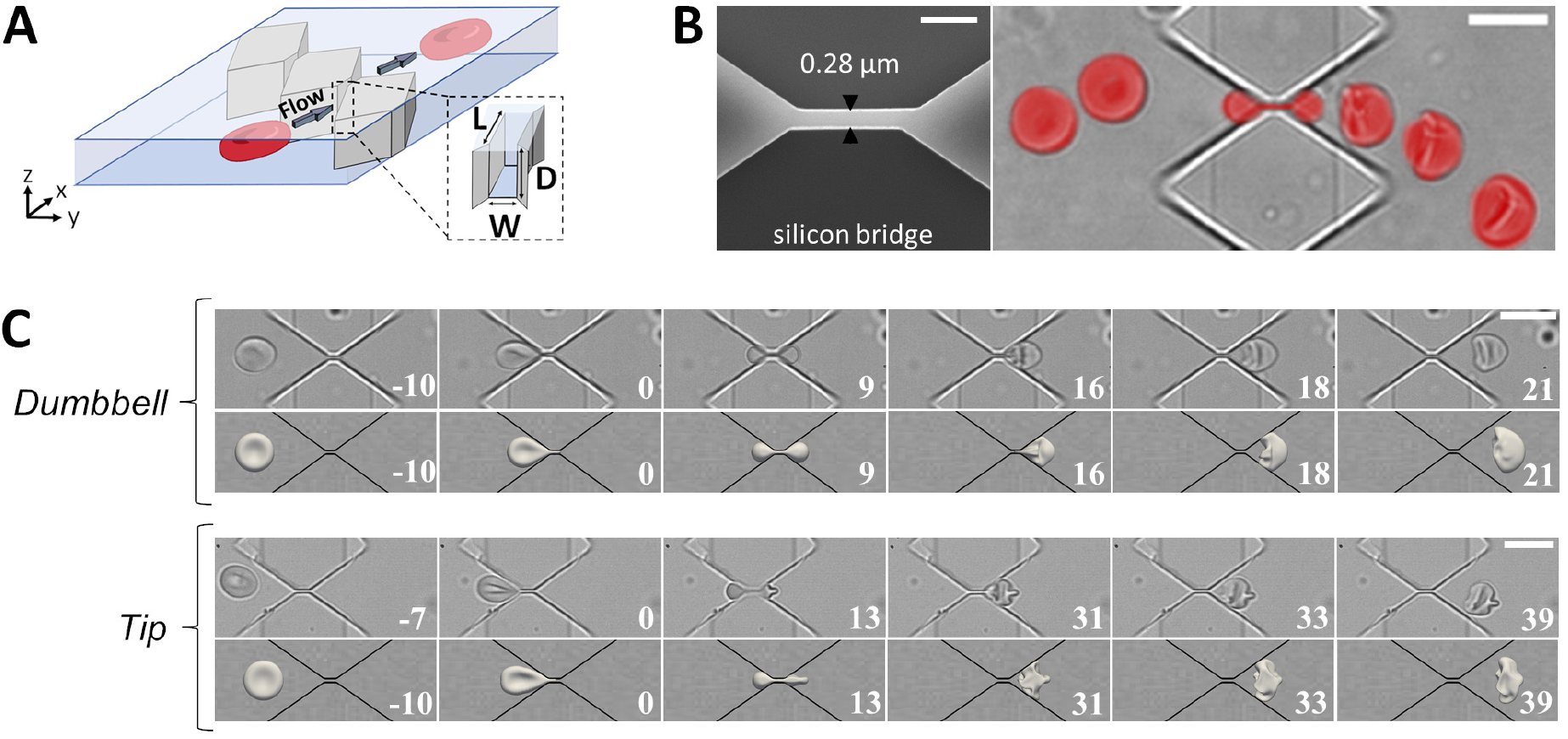
Principle of the experiment and typical behaviors of RBCs in submicron-wide slits. (A) Schematics of an RBC approaching a slit. The x, y, and z axes indicate the flow direction, width, and depth of the channel, respectively. (B) RBCs can pass through 0.28-µm wide slits. Left: Electron microscopy image of a 0.28-µm wide silicon bridge (in light grey) used for molding 0.28-µm wide slits; scale bar: 1 µm. Right: Superimposed images of an RBC (in red) passing through a 0.28*×*1.87*×*5.0 µm^3^ slit (bottom view in the XY plane); the images are extracted from Movie S1; scale bar: 10 µm. (C) Experimental observations (top rows) and numerical simulations (bottom rows) of transiting RBCs displaying dumbbell and tip shapes at slit exit, in 0.74*×*2.12*×*4.80 µm^3^ (top panel) and 0.86*×*2.75*×*4.70 µm^3^ (bottom panel) slits, respectively. Time in ms. Scale bars: 10 µm.

### RBCs pass through 0.28-µm wide slits at 37 °C

We first aimed to determine whether RBCs could pass through the narrowest slits measured by electron microscopy. We prepared slits of 0.28 µm in width (*L* = 1.9 µm, *D* = 5.0 µm) and we observed that almost 100% of RBCs successfully passed through these slits at 37°C under a physiological pressure of 490 Pa. Even under an in-slit pressure drop as low as 190 Pa, no retention was observed in a slit of 0.37 µm (*L* = 1.4 µm and *D* = 5.4 µm) that we will consider in the following as the ‘reference slit’. However, at room temperature, retention was observed in slits of width less than 0.5 µm. Typically, 20% of cells are retained in a slit of 0.42 µm in width (*L* = 2.2 µm, *D* = 5.4 µm) under an in-slit pressure drop of 400 Pa (see *SI Appendix*, Table S1).

In addition to the impressive narrowness of the slits that RBCs are able to pass through, it is interesting to note the difference between the retention rate of RBCs observed at room temperature and at 37°C. As retention of an RBC results from its too low ability to change its shape under an applied pressure, this suggests that there is a thermally activated mechanism responsible for an increase of RBC deformability at 37°C.

### Mechanisms of retention/passage and deformation of RBCs

It is well established (25) that, under an external force, cell deformation occurs primarily by two mechanisms. The first mechanism is the redistribution of its volume enclosed in the inextensible lipid membrane, at constant surface area *S*, and volume *V*. The higher the sphericity index of the RBC, defined as *SI* = *π*^1*/*3^(6*V*)^2*/*3^*/S*, the lower its ability to deform. Indeed, a perfect sphere of *SI* = 1 cannot deform at constant surface area and volume. The second mechanism is the local resistance of the elastic spectrin network to shear and stretch, and the resistance of the membrane to adopt acute curvatures, so that not all geometric shapes compatible with a given surface area-to-volume ratio are accessible to the cell. The accessible shapes depend on the applied mechanical load.

### Geometrical aspects: the surface area of an RBC allows it to deform into two tether-connected equal spheres

As Canham and Burton (26) pointed out years ago, the purely geometric conditions for an RBC to pass through a slit of any width, length, and depth are that the cell has enough surface area to be able to form two spheres on either side of the slit opening, connected by an “infinitely” thin tether (Fig. 2A). It is when the cell is equally divided into two equal spheres on both sides of the slit that the surface area is maximum for a given cell volume. This configuration is the one with the highest sphericity index allowing the passage of an RBC through a slit, regardless of the slit dimensions. We name it two tether-connected equal sphere model, or two equal-sphere model in this study. A simple calculation of the relation between the surface area *S* and the volume *V* of two equal spheres gives the relation *S* = (72*πV* ^2^)^1*/*3^ = 6.09 *V* ^2*/*3^ with a sphericity index *SI* = 0.7937. This shape is however unrealistic. It ignores the membrane/cytoskeleton rigidity, which resists to curvature, shear and stretch in the tether, and the non-zero thickness of the tether, which consists of two membrane bilayers and the inner hemoglobin solution. These points will be discussed in the next section. Nevertheless, the effective cell deformation observed experimentally and numerically (Fig. 1 and Fig. 2B-E) shows the extreme ability of RBCs to deform and approach the two tether-connected equal sphere shape. Indeed, as seen in Fig. 2B (bottom view of an RBC in a slit), the projections of the rear and front parts of the RBC, upstream and downstream of the slit respectively, form an arc of a circle whose radius is equal to half the slit depth (Fig. 2C). This shows that the RBC has a spherical cap upstream and downstream of the slit. In addition, our numerical simulations that visualize the shape of the RBC inside the slit clearly show that the cell forms a narrow neck that does not fill the entire slit’s depth (Fig. 2D). Confocal imaging of RBCs in slits confirms the existence of this neck between the two RBC parts on both sides of the slit (Fig. 2E and *SI Appendix*, Movie S4).

**Fig. 2.**
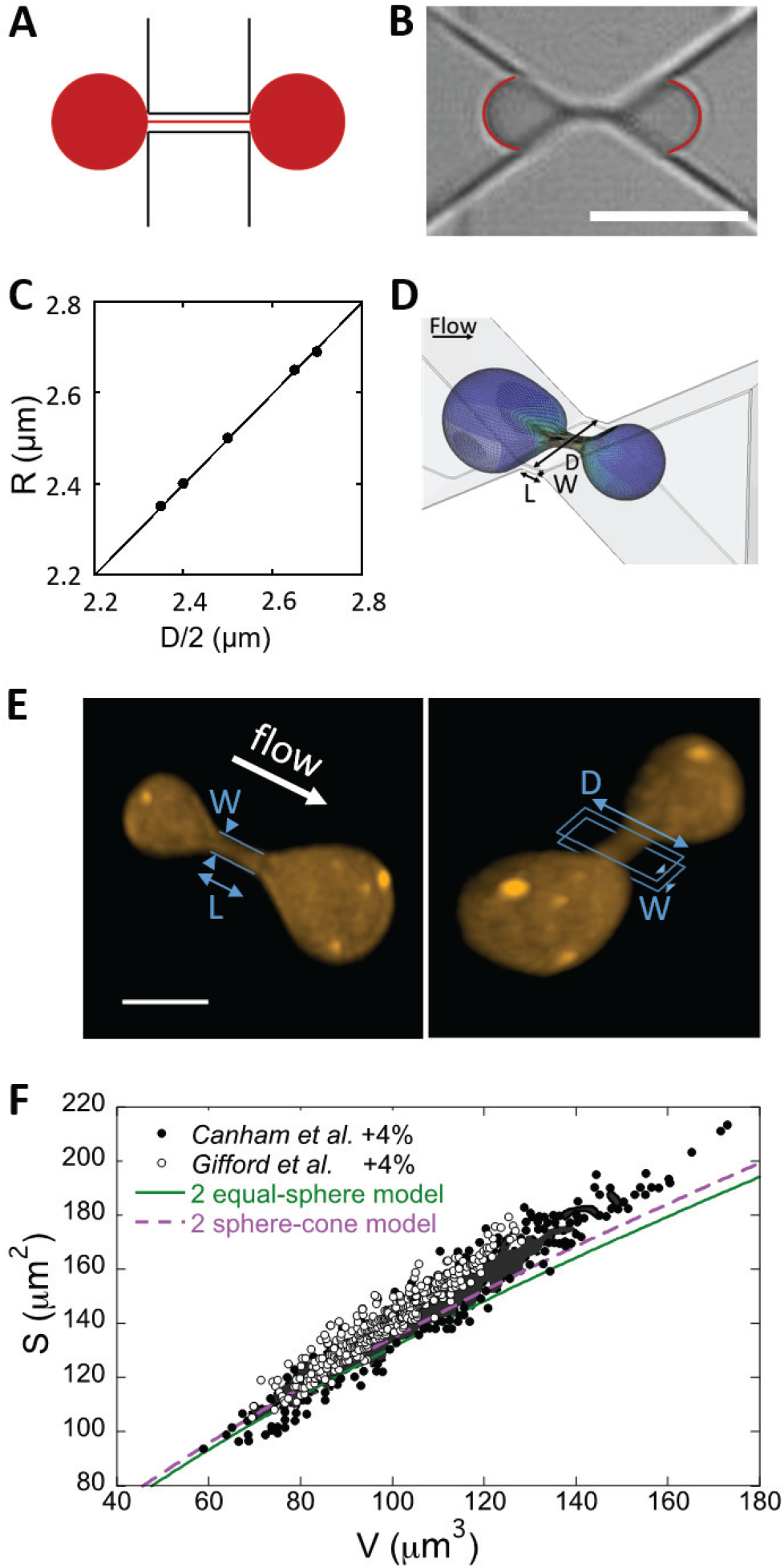
RBC shape in a slit. (A) Schematics of two equal spheres connected by an infinitely thin tether. (B) Image of an RBC symmetrically positioned in a 0.37*×*1.33*×*4.7 µm slit at 37°C, with two arcs of circle superimposed at its front and rear. Scale bar: 5 µm. (C) Radius of front and rear arcs of circle averaged on 10 RBCs symmetrically positioned in slits versus half slit depth *D/*2; temperature from 17°C to 37°C; in-slit pressure drop Δ*P* = 500 Pa. The solid line is the bisector. (D) Simulation of an RBC passing through a 0.28*×*1.9*×*5.0 µm^3^ slit at 37°C under an in-slit pressure drop 500 Pa. Inside the slit is formed a thin neck that does not fill the slit’s depth. (E) Top and tilted views of the 3D-reconstruction of an RBC in a 0.67*×*2.34*×*4.98 µm^3^ slit. The slit walls are drawn in blue as a guide to the eyes. Scale bar: 3 µm. (F) Surface area *S* versus volume *V* of circulating RBCs corrected from Refs. (26, 27), two equal-sphere curve (solid green line) and approximate two cone-sphere curve mimicking the experimental slit geometry (dashed purple line).

This geometric ability, characterized by *SI* ≤ 0.7937, was tested on data of surface areas and volumes measured at room temperature on RBCs collected from the microcirculation in two previous studies (26, 27). Figure 2F shows the surface area-volume relation of thousands of circulating RBCs together with the *S* = 6.09 *V*^2/3^ curve corresponding to the two equal-sphere shape (*SI* = 0.7937). To account for the fact that RBCs are at 37°C in the spleen, their surface area was first increased by 2%, which is the surface dilation of the cell membrane at 37°C (2). In a second step, it was further increased by 2% to account for the ability of the cell membrane to expand slightly under tension without breaking (2). It is striking to note that almost all cells have, for a given volume, a surface area at least equal or higher than that required to form two equal spheres and a tether (green curve in Fig. 2F). Even more striking is the fact that experimental data are very close to the two equal-sphere curve, even though they deviate from the curve for the largest cells. This strongly suggests that it is the mandatory passage through the splenic slits that determines the surface area-to-volume ratio of circulating RBCs.

It is worth noting that our slits have oblique walls. Figures 2B and D show that the shape of the RBC parts upstream and downstream of the slit is not spherical but roughly looks like a cone of revolution ended by a spherical cap. This thus indicates that the RBC surface area is slightly higher than that required in the two equal-sphere model. By approximating the RBC shape in the slit as two cones of revolution ended by spherical caps and connected by an infinitely thin tether, the *S* versus *V* relationship becomes *S* = 6.25 *V*^2/3^ (see the derivation in *SI Appendix*, Section 3). The corresponding curve (dashed purple line in Fig. 2F) clearly shows that most of the circulating cells from Refs. (26, 27) still have a surface area-to-volume ratio allowing a deformation into a two cone-sphere shape, in agreement with our observations. The crucial role of RBC morphology is illustrated by the behavior of a spherocytic RBC and a highly elongated irreversible sickle RBC from a patient with sickle cell disease. The spherocyte has no surface area reservoir. It is therefore unable to deform in a two-sphere shape and is blocked at the entrance to the slit (*SI Appendix*, Movie S5). Conversely, the irreversible sickle RBC successfully passes through the slit (*SI Appendix*, Movie S6) as the cytoplasmic hemoglobin fibers probably aligns along the flow direction and its overall shape is sufficiently elongated to pass through the slit without further deformation. Obviously, however, the actual strain achieved by an RBC subjected to a given in-slit pressure drop, and in particular, its ability to form a thin tether within the slit, is limited by the cell rigidity. This actual strain determines the threshold dimension of the slit that the cell can pass through.

### Mechanical aspects: spectrin unfolding and membrane deformation

To have a deeper insight into the involved mechanical processes of deformation, we numerically calculated using ABAQUS software the minimum area, *S*_*m*_, required for an RBC of given volume *V* to pass through the reference slit under an in-slit pressure drop of 500 Pa. The mechanical parameters taken into account in the calculation are the shear modulus of the cytoskeleton *µ* (a function of deformation as shown in Eq. (2) in *SI appendix*, Section 2, *µ*0 is the initial value of *µ* without any deformation), the area compression modulus of the bilayer *K*, and the force necessary to unfold half of the spectrin domains of a spectrin tetramer *F*_1*/*2_ (*F*_1*/*2_ = 10 pN and *K* = 450 mN/m at 25°C; *F*_1*/*2_ = 1 pN and *K* = 375 mN/m at 37°C) (2, 28–31). Regarding the *F*_1*/*2_ force, the meshes of the spectrin network can indeed distend very significantly when a mechanical stress is applied to the RBC membrane (28). This phenomenon is due to the unfolding of domains in spectrin molecules subjected to an external force, whose minimum value required for unfolding depends strongly on the temperature (29). The computed variations of *S*_*m*_ with *V* are shown in Fig. 3A-B, together with the experimental data from Gifford et al. (27) corrected to account for the possible 2%-membrane area extension without lysis, and, at 37°C, 2%-thermal dilation of the lipid bilayer. We focus on Gifford’s data as the microfluidic technique to extract *S* and *V* is considered more accurate. At 25°C, when *F*_1*/*2_ and *K* are set to 10 pN and 450 mN/m, respectively, (32), we clearly observe in Fig. 3A that the numerical curve *S*_*m*_(*V*) (blue solid line) significantly cuts the experimental *S*(*V*) curve. On the other hand, at 37°C (Fig. 3B), when *F*_1*/*2_ and *K* are set to 1 pN and 375 mN/m, respectively, the vast majority of the (*S,V*)-experimental data are above the numerical curve (red solid line). To estimate the RBC population that can pass through a slit without needing to unfold spectrin domains, we applied the previous calculation for a very high unfolding energy, inaccessible to the cell, i.e. *F*_1*/*2_ = 100 pN. The corresponding *S*_*m*_(*V*) curves are shown in dashed blue at 25°C and dashed red at 37°C. Above the dashed curves, RBCs have therefore sufficient surface area to pass through the slits without unfolding spectrin domains. Between solid and dashed curves, RBCs must locally unfold spectrin domains, mainly inside the slit when the cell forms a neck (grey zones in Fig. 3C) with the conditions *F*_1*/*2_ = 10 pN at 25°C and *F*_1*/*2_ = 1 pN at 37°C). At 25°C (Figure 3A), 50% of RBCs are retained, 11% pass with unfolded spectrin domains and 39% pass without spectrin unfolding. Spectrin unfolding therefore rarely occurs a 25°C due to the high value of *F*_1*/*2_, while at 37°C (Figure 3B), 28% pass with unfolded spectrin domains, 62% pass without spectrin unfolding and only 10% of RBCs are retained. In another word, if no spectrin unfolding was allowed, then the retention rate would be 38% rather than 10% at 37°C. It is worth noting that the RBC percentage requiring spectrin domains unfolding depends on slit dimensions, in-slit pressure drop and temperature. Both simulated behaviors are in full agreement with our experimental observations of retention in the reference slit at both room temperature and 37°C. In Fig. 3D, the minimal pressure drop Δ*P*_*c*_ required for an RBC to pass through the reference slit is calculated and plotted versus the cell sphericity index *SI*. Interestingly, we observe that RBCs of *SI* < 0.768 are able to pass through the reference slit under a physiological pressure Δ*P* = 500 Pa (24) although the RBCs must be much floppier to pass through it at 25°C (*SI* < 0.75). Figure 3E illustrates the critical pressure drop required for an RBC with a standard shape to pass through a slit of length and depth equal to those of the reference slit as a function of the slit width. Although the critical pressure drops are similar for large slits at both 25°C and 37°C, they are much higher at 25°C than at 37°C for the narrowest slits. It should be noted that the experimental points linearly extrapolated from the dynamic data described below are well within the range of values of the numerical predictions, which further strengthens our numerical approach. The simulations show that the differences of behaviors observed between the two temperatures are mainly due to the *F*_1*/*2_ contribution rather than bilayer area modulus K as shown in *SI Appendix*, Fig. S1. Our in-silico approach suggests that the cell to unfold spectrin domains is regurlarly tested by interendothelial slits.

**Fig. 3.**
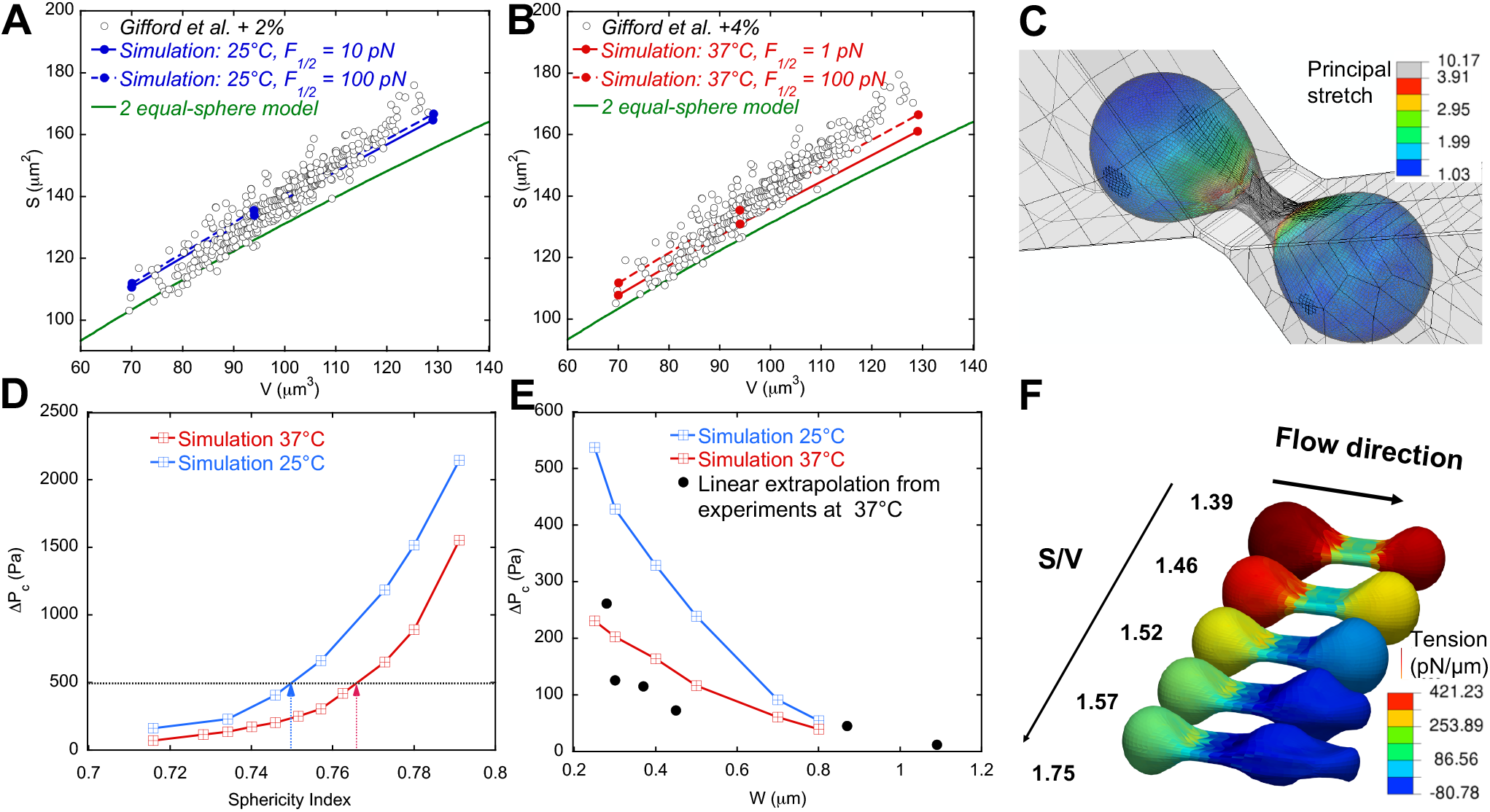
In-slit mechanisms of RBC deformation. (A-B) RBC corrected surface area from Ref. (27) (open circles) and computed minimal surface area required to transit through a 0.38*×*1.3*×*4.7 µm^3^ slit at Δ*P* = 500 Pa (solid and dashed lines) versus RBC volume at 25°C (A, blue lines) and 37°C (B, red lines), with the two equal-sphere model (green solid line). (C) RBC deformation (principal stretch in color scale) and spectrin unfolding (region with spectrin unfolding is marked in grey), calculated in ABAQUS computation. The volume and surface area of the RBC are 97.9 µm^3^ and 135.1 µm^2^ which is close to the red solid curve in B. The slit dimensions are 0.38*×*1.3*×*4.7 µm^3^, and temperature is 37°C. (D) Critical pressure drop Δ*P*_*c*_ required to pass through the reference slit versus sphericity index at 25°C and 37°C. The black dashed line represents the average physiological pressure of 500 Pa, and the blue and red arrows represent the maximal sphericity index needed to pass through under this physiological pressure. (E) Critical pressure drop Δ*P*_*c*_ versus slit width *W* (*L* = 1.3 µm, *D* = 4.7 µm) for the RBC standard shape. Experimental data were obtained from measurements performed at 37 °C of the RBC transit time (*t*_*t*_) through slits of varying width under different applied pressure drops, by linearly extrapolating the curves 1/*t*_*t*_ versus Δ*P* at 1/*t*_*t*_ = 0 (see Fig. 5A). (F) RBC shapes and surface tension (color scale) in a 0.80*×*2.0*×*5.0 µm^3^ slit under an in-slit pressure drop of 350 Pa for various surface area-to-volume ratios (unit: 1/*µ*m).

Next, we tested the role of membrane rigidity by treating RBCs with 1 mM diamide, which increases the shear modulus and viscosity of the cell membrane (33, 34), and observing their passage through 0.61*×* 2.01*×* 5.0 µm^3^ slits. No major change was observed in the retention rate, which remained below 10% (see *SI Appendix*, Table S1). This result is in line with those reported on rat studies (with sinusoidal spleen like human), ex-vivo isolated-perfused human study (34) and clinical observations on Southeast Asian ovalocytosis patients (35), which indicates that decrease in membrane deformability does not strongly affect the RBC ability to cross the splenic slits. We also conducted numerical simulations to predict the effects of increased membrane viscoelasticity due to diamide treatment. For the control case, the membrane viscoelasticity is modeled using a multiscale model described in *SI Appendix*, giving an initial shear modulus of the cytoskeleton *µ*_0_ = 9 pN/µm. The characteristic time *t*_*c*_ = *η*_*m*_*/µ*_0_ is chosen as 0.1 s as measured in experiments (2), where *η*_*m*_ = 0.9 pN*·*s/µm is the membrane viscosity. For the diamide-treated case, we increased both the initial shear modulus and membrane viscosity by 10 times (33, 34) while keeping *t*_*c*_, *F*_1*/*2_, and *K* unchanged. We found that the critical pressure for the cell to pass the slit is almost the same in the control and diamide-treated cases, which is consistent with our previous numerical study (18) and experimental studies by others (34). Finally, we characterized the role of *SI* on the shape of transiting RBCs. Figure 3F shows the local membrane tension of an RBC of volume 94 µm^3^ at a given position in a 0.8 *×*2.0*×* 5.0 µm^3^ slit as it passes through under an in-slit pressure drop of 350 Pa. The surface area-to-volume ratio was varied from 1.39/*µ*m to 1.75/*µ*m (*SI* from 0.776 to 0.617) by increasing the surface area while keeping a fixed volume. It clearly shows that the membrane tension is maximal in both RBC bulges when *SI* is high, but when *SI* is lower than 0.74 (surface area-to-volume ratio of 1.44/*µ*m), the membrane tension in the downstream part of the cell is low. For the most deflated cells, we even see that the membrane tension is negative in the downstream part, which induces a curvature inversion in the front of the cell, generating tips that are observed experimentally (Fig. 1C). It is worth noting that the membrane tension range does not exceed a few hundreds mN/mm, 20 times below the RBC lysis tension (2).

### No calcium-related active process is detected

It is now known that the RBC membrane contains a mechanosensitive ion channel, PIEZO1, which opens upon cell membrane stretching, inducing an internal influx of calcium from the cell environment. In turn, this influx activates the Gardos ion channel which induces an outflow of potassium and water from the cell (36). The combination of PIEZO1 and Gardos could therefore promote the deflation of RBCs (36–38) and play a role in their passage through the slits. Note that this activation is not necessary to pass through 0.28 µm slits since the RBC suspension buffer used in the previous sections did not contain calcium. We quantified the change in intracellular free calcium in RBCs at the exit of 0.8-µm wide slits in the presence of 1mM calcium in the RBC suspending buffer (Fig. 4). The amount of free calcium in RBCs significantly increases at the slit exit under in-slit pressure drops of 100 Pa (Fig. 4B), 500 Pa, and 1000 Pa (*SI Appendix*, Fig. S2). Interestingly, this behavior is observed both in the absence and presence of calcium in the suspension buffer and is therefore not specifically due to PIEZO1 activation. It might be attributed to a release of intracellular calcium, initially chelated by calmodulin (39). The calcium signal increases by a factor of 1.5 to 3 at the slit exit, both with and without calcium in the suspension buffer. Note that based on a physiological intracellular calcium concentration of 0.1 µM (40), the increase in calcium is still less than the 0.5 µM concentration needed to activate Gardos (41). In conclusion, no specific calcium entry that could be attributed to PIEZO1 is observed during the RBC passage through the slits. One reason could be that PIEZO1-mediated calcium entry occurs within seconds, as suggested in Ref. (36), which is much longer than the transit times measured here (about 100 ms).

**Fig. 4.**
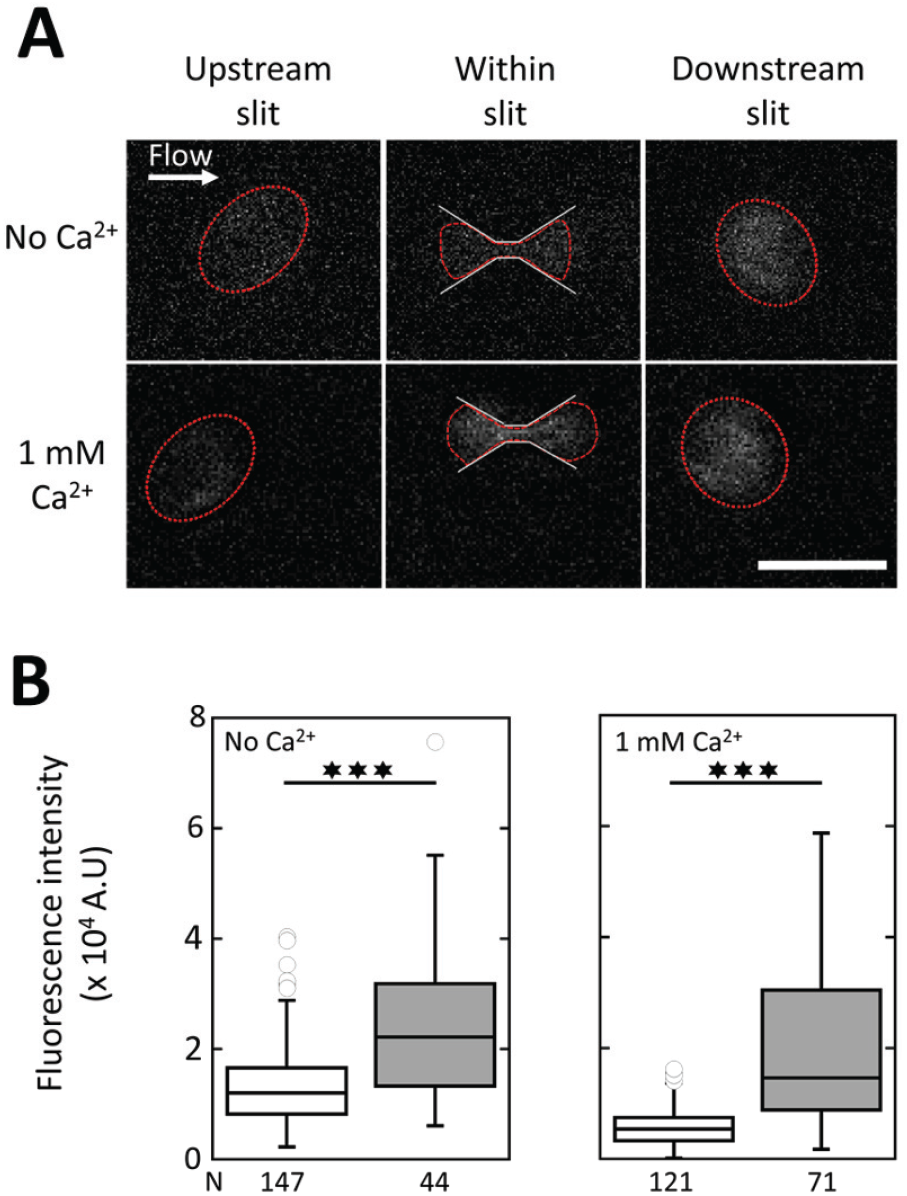
No calcium influx is detected during RBC passage through the slits. (A) Epifluorescence images of intracellular calcium in RBCs upstream of, within, and downstream of 0.80*×*2.77*×*4.70 µm^3^ slits, in absence (top row) and presence (bottom row) of 1 mM calcium in the external medium. The RBC contours are drawn in red. The white arrow indicates the flow direction. In-slit pressure drop ∆*P* = 490 Pa. Scale bar: 10 µm. (B) Intracellular calcium fluorescence intensity upsptream (in white) and dowsntream of the slits (in grey) in absence (left) and presence (right) of calcium under Δ*P* = 100 Pa. Median values are displayed with 25% and 75% percentiles, and min/max values as whiskers. N: number of analyzed RBCs.

### Dynamics of transit

Transit times are fast, sub-second, from 0.46 s in the 0.28 µm-wide slit under 500 Pa to 1.9 ms in the widest slits (*W* = 1.09 and 0.86 µm under 660 Pa and 1200 Pa, respectively) (see *SI Appendix*, Table S1 for a summary of measured transit times). These values, observed under in-slit pressure drops of 500 Pa or less, including rare long events, are in the range of those observed by intravital microscopic videorecordings of Ringer-perfused mouse spleens (42). For each slit size, a few rare long events are observed (*SI Appendix*, Fig. S3). The variation of the inverse of the median transit time, 1/*t*_*t*_, i.e. a quantity that is an effective velocity across the slit, varies linearly with the in-slit pressure drop in the range of 200 Pa to 2000 Pa for slit widths from 0.28 to 1.09 µm and at the three temperatures we studied, 15°C, 22°C, and 37°C (Fig. 5A for 37°C and *SI Appendix*, Figs. S4 and S5 for 22°C and 15°C, respectively). Numerical simulations of transit times were performed in the same range of pressure drops and temperatures and for slits wider than 0.6 µm to ensure reasonable simulation times. The quantitative agreement between experimental and numerical values is excellent for all slit dimensions, pressures and temperatures (Fig. 5A and *SI Appendix*, Figs. S4 and S5). Note that the viscosity of the inner hemoglobin solution used for the numerical simulations are 6.9, 9.7, and 17 mPa.s, at 37°C, 22°C, and 15°C, respectively. These values are in the range of those recently found by using molecular rotors as intracellular probes of RBCs (43). The transit times of diamide-treated RBCs were explored under different pressure drops. The RBC passage is clearly slowed down (see *SI Appendix*, Fig. S6), being the signature of an increase in viscosity, likely in membrane viscosity as diamide is known to not increase the cytoplasmic viscosity (33, 44). Indeed, numerical simulations fit well the experimental data by using an initial cytoskeleton shear modulus *µ*0 = 90 pN/µm and a membrane viscosity *η*_*m*_ = 9 pN. s/µm (*t*_*c*_ = 0.1 s), which are 10 times those of the control case without diamide (*SI Appendix*, Fig. S6), while increasing *µ*0 alone didn’t slow down the passage. It is interesting to note that our results strongly suggest that diamide treatment not only alters membrane shear modulus but also membrane viscosity, which is consistent with the existing studies (33, 44). The role of the cytoplasmic viscosity was also explored by performing experiments with RBCs suspended in hypertonic solution (400 and 500 mOsm) to decrease their volume in order to increase their inner hemoglobin concentration and, by consequence, their cytoplasmic viscosity. The two factors (volume and cytoplasmic viscosity) act in opposite ways on cell transit time. On the one hand, a smaller RBC volume passes through the slit faster. This tends to shorten the transit time. On the other hand, the increased cytoplasmic viscosity results in a longer transit time. We show in *SI Appendix*, Fig. S7 that the viscosity effect dominates because the transit time of cells suspended in a hyper-osmotic medium increases. We also carried out a parametric study on how quantitatively the transit time changes with the increased mean corpuscular hemoglobin concentration (MCHC) and decreased volume under hypertonic conditions using numerical simulations. The increased hemoglobin solution viscosity *η*_*Hb*_ with elevated MCHC was calculated using the Ross and Minton model 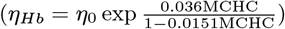(45) based on experimental data (see (*SI Appendix*, Fig. S8), where *η*0 = 0.7 mPa·s is the water solvent viscosity at body temperature. From these simulations, we estimated the MCHC that leads to experimentally measured transit time (*SI Appendix*, Fig. S8). For example, the MCHCs at 400 and 500 mOsm were estimated as 38.9 and 42.2 g/dL, which are consistent with existing measurements of MCHC at different osmolarities (46).

**Fig. 5.**
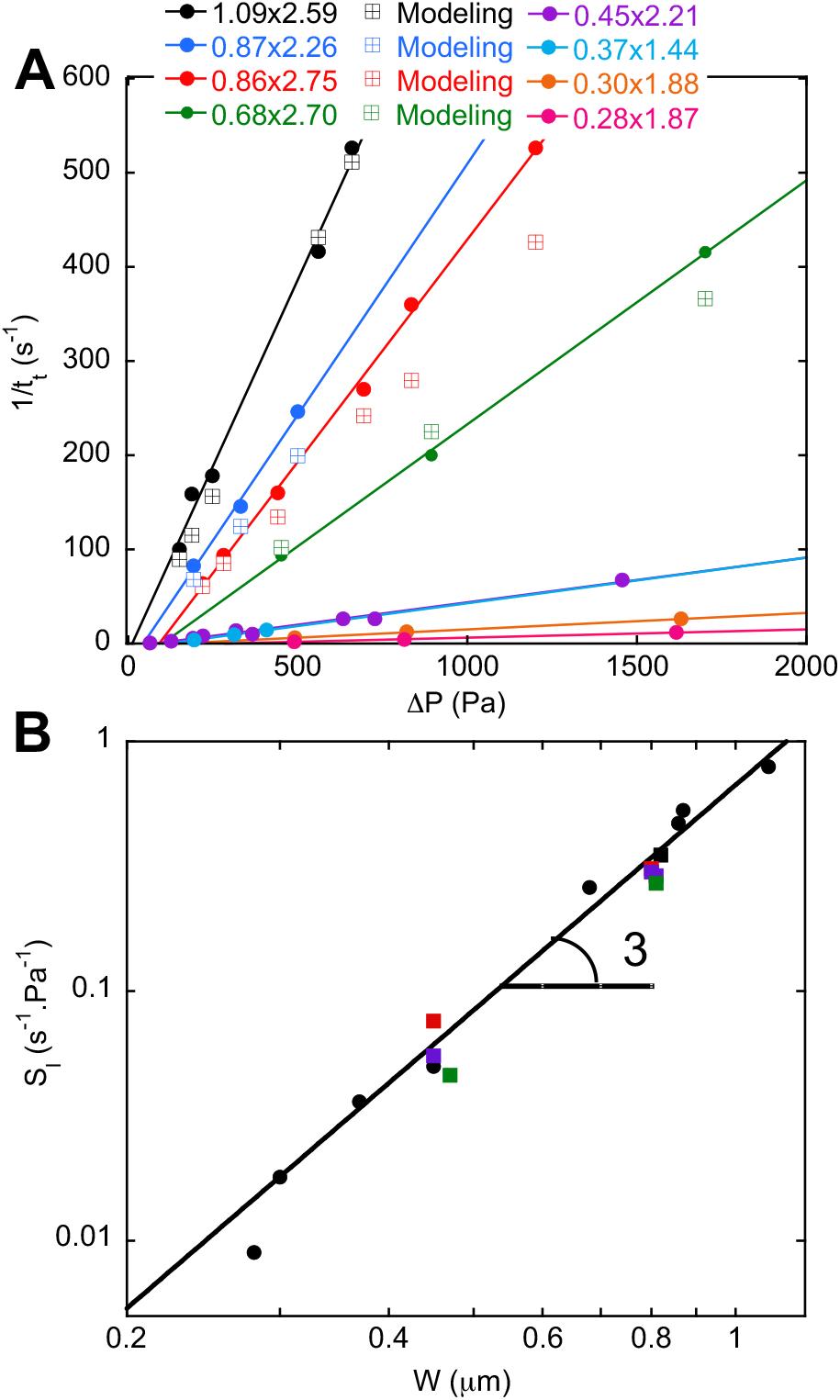
Dynamics of RBC passage through the slits. (A) Inverse of transit time 1*/t*_*t*_ versus in-slit pressure drop ∆*P* at 37°C for slits of different dimensions, from 1.09 to 0.28 µm in width: experimental data (closed circles), simulations (open squares with cross). The lines are linear fits of the experimental data. (B) Log-log representation of the 1*/t*_*t*_(Δ*P −* Δ*P*_*c*_) slope *S*_*l*_ versus slit width at 37°C. Black closed circles: no calcium, no plasma; black closed squares: no calcium, plasma; red closed squares: 1 mM calcium, plasma; purple closed squares: 1 mM calcium, plasma, 5 µM GsMTX4: green closed squares: 1 mM calcium, plasma, 10 µM TRAM34. The solid line is a fit of *S*_*l*_ versus *W* ^3^

To test the possible role of PIEZO1 and Gardos channel activation on the dynamics of RBCs in slits, experiments were also conducted in the presence of 1 mM calcium in the suspension buffer and then adding 5 µM of the PIEZO1 inhibitor, GsMTX4, or 10 µM of the Gardos inhibitor, TRAM34. We observed a linear behavior of the inverse of the transit time with applied pressure drop similar to the control conditions without inhibitors, as shown for two slit sizes (*SI Appendix*, Fig. S9). Thus, in our experimental conditions, we do not see any significant impact of the channels’ inhibitors on the transit times.

From the linear fits of 1*/t*_*t*_ versus Δ*P* (Fig. 5A), two parameters were extracted. The first one is a dynamical critical pressure drop that we take as the value of the pressure drop of the fitted line extrapolated at 1/*t*_*t*_ = 0. This value gives only an order of magnitude of the critical pressure drop required for an RBC to pass through the considered slit because i) this value is extrapolated from the median values of the transit times at a given pressure and, as we showed in Fig. 3D from ABAQUS computations, the critical pressure drop strongly depends on *SI* of individual RBCs, and ii) the relationship between 1/*t*_*t*_ and Δ*P* is probably non-linear at low Δ*P* values when the effect of membrane stress becomes more important. Despite these limitations, the values of the dynamical critical pressure drops we extrapolated from 1/*t*_*t*_ versus Δ*P* variations are in the same range as those calculated from ABAQUS computations (Fig. 3E). This confirms the force of our coupled experimental/numerical approach. The second parameter we extracted is the slope of the 1/*t*_*t*_ versus Δ*P* linear variations. This slope, 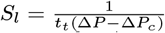is plotted in Fig. 5B versus the slit width *W* in a log-log representation. The *S*_*l*_ versus *W* variations are well fitted by a power law of exponent 3. An identical result is obtained at 22°C and 15°C (*SI Appendix*, Fig. S10). It shows the very strong dependence of the RBC dynamics to the slit width.

It is interesting to note that the dynamic behavior of RBCs is similar to that of a 2D Poiseuille flow with a Newtonian fluid (47). Let’s consider an RBC with a volume *V* and interior viscosity *η*_*d*_ passing from the left to the right through a slit of width *W* and depth *D*. To derive a simplified closed form estimation of the transit time *tt* of this process, we consider it as a 2D Poiseuille flow with a depth of *D*. The pressure drop is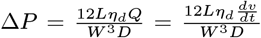, where *Q* is the flow rate and *v* is the volume of the RBC on the right side of the slit, which is almost zero in the beginning and becomes *V* in the end of the process. Integrating the equation as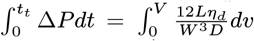we get 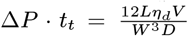. This relationship is in excellent agreement with our experimental observations of the dependence of transit time *t*_*t*_ on Δ*P* (Fig. 5A) and *W* (Fig. 5B), although in this simplified analysis we ignored the effect of membrane stress, the pressure drop of the flow entering the slit (Sampson/Roscoe flow (48)), the short time for an RBC to enter and leave the slit in the beginning and the end of the process, and the contribution from the much less viscous exterior fluid. This agreement strongly indicates that it is the inner hemoglobin solution that mainly governs the RBC dynamics through a slit of given dimension. Finally, Fig. 5B clearly shows that the passage dynamics of RBCs through the slits is not affected either by the presence of calcium in the suspension or by the inhibitors of PIEZO1 or Gardos channels. These results reinforce the idea that potential activation of PIEZO1 in the slits does not play a significant role in the passage of RBCs through the splenic slits, either in terms of retention or passage dynamics.

## Discussion and conclusion

The first splenic selection of RBC deformability is of geometrical origin. The strong dumbbell cell deformation with a very thin neck inside the slit and spherical caps at the entrance and exit of the slit that we evidenced indeed requires that the RBC surface area-to-volume ratio be higher than that of two equal spheres. This is observed, strikingly, on circulating RBCs. Even more striking is that the surface area-to-volume ratio of circulating RBCs is not more than 15% larger than that of two equal spheres. Thus, there is no additional benefit to having more excess surface area than that required to pass through the slits. Yet, a greater excess of surface area would decrease the membrane tension and limit the need for spectrin unfolding in the slits. This is not the case retained by Nature, maybe because an excess of surface area has a high biological cost or is quickly lost by membrane vesiculation as RBCs pass through the slits. The second splenic selection of RBC deformability is of mechanical origin. We targeted the two identified pathways potentially able to dynamically modify RBC mechanical properties. The first pathway involves the mechanosensitive PIEZO1 channel known to participate in regulating RBC volume. Our experimental results strongly suggest that no PIEZO1 effect is involved in the RBC passage. This may be due to the too short transit times of RBCs through the slits that do not allow a significant calcium influx. Indeed, most RBC transit times were less than one hundred milliseconds under physiological conditions in agreement with those previously observed for mice (42). The second pathway involves the spectrin network, which underlies the RBC membrane and is known to unfold under an external mechanical stress. We used an in-silico approach, as to our knowledge there is no direct method for visualizing spectrin unfolding in non-modified living RBCs (28). It shows that local spectrin unfolding within the narrowest slits is crucial for the passage of RBCs with less surface area. We emphasize that our coupled experimental/simulation approach is quantitative and can address the critical need for in-vitro assessment of splenic clearance of diseased or modified RBCs for transfusion and drug delivery. In addition, our approach provides a basis for further developments, including the study of RBC pitting, an important function of the spleen where vacuoles, bodies, or even malaria-causing parasites internalized in RBCs are expelled from the host cell during slit passage. Finally, the enucleation of the RBC erythroid precursor during bone marrow egress could also be explored using the approach presented here.

## Materials and Methods

More details are available in *SI Appendix*.

### Blood samples and buffers

Human RBCs were obtained from voluntary healthy donors who were fully informed regarding the purposes of the study and gave their consent. All blood samples were de-identified prior to use in the study. RBCs were used within 6 hrs, except for spherocytic and sickle RBCs which were delivered 1 day after harvesting from patients. RBCs were stored in DPBS-G (Dulbecco’s Phosphate Buffered Saline solution (Gibco) + glucose) at a volume fraction (hematocrit, Hct) of 50%, and diluted at 1% Hct right before experiment. When calcium was present in the external medium, DPBS-G was supplemented with 10% plasma. Membrane rigidity was increased with 1 mM diamide (Sigma-Aldrich). PIEZO1 and Gardos ion channels were inhibited with 5 µM GsMTx-4 (Alomone Labs) and 10 µM TRAM34 (Sigma-Aldrich), respectively. RBC membrane and intracellular calcium were labelled using CellTrace™ Yellow and Fluo-4 AM (ThermoFisher), respectively.

### Microfluidic device fabrication

The microfluidic device was made of polydimethysiloxane (PDMS, Sylgard 184, Dow Corning) and consisted of a PDMS chip, molded from a silicon master, and covalently assembled to a PDMS-coated glass coverslip. The silicon master mold was fabricated following a protocol adapted from the procedure described in Ref. (49).

### Flow experiments and microscopy

The microfluidic device was placed on an inverted microscope (Olympus IX71) equipped with 20× /60× /100*×* objectives, temperature controller, and cameras. The device surfaces were coated with 1% BSA before injection of 1% Hct suspensions. The flow of the RBC suspension was pressure-controlled (MFCS-8C, Fluigent). Typically, RBCs were observed in brightfield microscopy and movies were acquired at high frame rate using a high-speed camera (Fastcam Mini camera, Photron). For measurement of intracellular calcium, RBCs were observed at low frame rate in epifluorescence with the 100*×* objective using an sCMOS camera (Neo 5.5, Andor). 3D-imaging of RBCs in slits was performed using a benchtop confocal microscope equipped with a 60× objective (BC43, Andor). For each microfluidic chip and each experiment, streamlines, pressure drop, and fluid velocity in the slits were calculated using COMSOL® software (see *SI Appendix*, Fig. S11).

### Video analysis

Analysis of RBC retention was done using FIJI software (ImageJ): retained RBCs were counted with respect to the total of retained and passing RBCs. The retention rate was considered as significant for a percentage of retained RBCs higher than 10% in a population of at least 50 RBCs. Analysis of RBC transit through slits was carried out either manually (FIJI) or using a custom Matlab routine: the transit time was measured as the time spent by an RBC in contact with the slit. Analysis of RBC intracellular calcium was done using FIJI: the calcium fluorescence intensity was measured over the RBC projected area upstream of, within, and downstream of the slit.

### Dynamic and quasi-static simulations of RBCs passing through slits

For dynamic simulations, we applied an in-house code with coupled finite element (FEM) and boundary integral methods to study the dynamics of RBCs passing through the slits and their transit time with full fluid-structure interaction considered (50). We simulated the full fluid-cell interaction under prescribed pressure drop. The boundary integral method was used to simulate the surround Stokes flows, while the FEM was used to model the RBC lipid bilayer and cytoskeleton as two interacting shells. The same mesh of lipid bilayer was used in finite element and boundary integral simulations so that the velocity and stress jump conditions on the interface were satisfied. For quasi-static simulations, we applied the commercial FEM package ABAQUS (23) to study the retention of RBCs in the slits and the corresponding critical pressure drop without modeling the fluid flow explicitly, because the dynamics, such as viscosity, does not play a role in the critical pressure and the quasi-static approach is much more efficient.

### Multiscale model of RBCs

The constitutive law for the RBC cytoskeleton in the finite element model was obtained using a multiscale model of RBCs. The mechanical response of a RBC membrane involve mechanics at different length scales, ranging from dynamics in the whole cell level (in the micrometer scale) down to the local dynamics of the spectrin and its tension-induced structural remodeling such as domain unfolding of spectrins in the protein level (in the nanometer scale). To capture these phenomena together, we developed a multiscale approach including three models at different length scales and connect them by a sequential information-passing multiscale algorithm. This multiscale model was implemented in both the in-house code and a ABAQUS VUMAT user subroutine (23).

## Data Availability Statement

All data are included in the manuscript and *SI Appendix*.

## ACKNOWLEDGMENTS

The project leading to this publication has received funding from France 2030, the French Government program managed by the French National Research Agency (ANR-16-CONV-0001 and ANR-20-CE17-0024). We thank I. Oze-rov and F. Bedu from PLANETE microfabrication facility. We thank Pr. Pierre Buffet and his group for providing us spherocytic and irreversible sickle RBCs. Z.P. and H.L. acknowledge the supports from US National Science Foundation grants NSF CBET-1706436/1948347 and NSF DMS-1951526. E.H. belongs to the French Consortium AQV.

