## Supplemental Information for "Physical mechanisms of red blood cell splenic filtration"

PNAS
www.pnas.org

### Supplementary Information for

#### Physical mechanisms of red blood cell splenic filtration

**Alexis Moreau, François Yaya, Huije Lu, Anagha Surendranath, Anne Charrier, Benoît Dehapiot, Emmanuèle Helfer, Annie Viallat, Zhangli Peng**

**Corresponding to Annie Viallat, Zhangli Peng.**
****

##### This PDF file includes:

Supplementary text
Figs. S1 to S11
Tables S1 to S2
Captions for Movies S1 to S6
References for SI reference citations

##### Other supplementary materials for this manuscript include the following:

Movies S1 to S6

#### Supporting Information Text

#### 1. Experimental methods

##### 1.1 Buffers, RBC samples, treatments and labeling

*Buffers.* The RBC standard suspension medium, DPBS-G, consists of Dulbecco's Phosphate Buffered Saline solution (DPBS 1X, no $CaCl_2$, no $MgCl_2$, Sigma-Aldrich), adjusted to pH 7.40, and to an osmolarity of 300 ± 5 mOsm by adding glucose. For standard microfluidic experiments, DPBS-G was supplemented with 0.5% bovine serum albumin (BSA, Sigma-Aldrich). For experiments with calcium in the external medium, DPBS-G was supplemented with 1 mM $Ca^{2+}$ and 10% human plasma (from voluntary donor, stored at -20°C). For experiments with varying osmolarity of the external buffer, we diluted a concentrated DPBS solution (DPBS 10X, no $CaCl_2$, no $MgCl_2$, Sigma-Aldrich) to desired osmolarities. Osmolarities were measured using a freezing point osmometer (Gonotec OSMOMAT 030).

*RBC samples.* For RBCs from healthy donors, a 30-µL blood droplet was obtained via pinprick and diluted in 1 mL DPBS-G. The RBCs were then washed and centrifuged at 500 g at RT three times in DPBS-G, and finally re-suspended in DPBS-G at a Hct of 50% as stock suspension. Experiments were performed within 6 hrs after RBC harvesting, with dilution of the RBC suspension to 1% Hct in DPBS-G + BSA or in DPBS-G + BSA + $Ca^{2+}$ + plasma right before the experiment. For spherocytic and sickle RBCs, blood samples harvested from patients were shipped within a day, then processed like healthy samples.

*Intracellular calcium labelling.* RBC intracellular calcium was labelled by incubating the 50% Hct stock solution with 5 µM Fluo-4 AM (ThermoFisher Scientific) under agitation for 3 h at 4°C. After incubation, RBCs were washed three times as above, resuspended at 50% Hct in DPBS-G for a few hours storage, before dilution to 1% Hct for the experiment.

*Ion channel inhibition.* PIEZO1 and Gardos channel were inhibited by incubating the RBCs at 1% Hct with 5 µM GsMTx-4 (Alomone Labs) or 10 µM TRAM34 (Saigma-Aldrich), respectively, for 20 min. Experiments were performed in presence of the ion channel inhibitors.

*Diamide treatment.* RBCs were washed three times and incubated during 1 hour at 37 °C in DPBS supplemented with 1 mM diamide (Sigma-Aldrich), a concentration known to have a significant effect on RBC membrane rigidity. After incubation, RBCs were washed twice and resuspended to 1% Hct for the experiment.

##### 1.2 Fabrication of the microfluidic device

The PDMS microfluidic device consisted of a chamber made of a bottom PDMS-coated glass coverslip assembled with a PDMS chip containing the microfluidic channel with slits.

*Silicon master mold fabrication.* The master mold was fabricated following a protocol adapted from the procedure previously described in Gambhire et al. (1). Briefly, the fabrication consisted of 3 main steps: (i) standard photolithography to fabricate the largest elements of the mold (main channel, inlets/outlet), (ii) electron beam lithography to produce large hexagonal hollows in the main channel separated by bridges of a few µm width, and (iii) anisotropic chemical etching in a mixed solution of 20% KOH / 10% IPA at 70°C to enlarge the hexagonal hollows and thin the bridges down to submicron width. All steps were carried out in the clean room facility PLANETE (CT-PACA Micro- and Nanofabrication Platform, Marseille, France).

*Chip fabrication and device assembly.* PDMS was mixed with the curing agent in a 10:1 ratio and poured on the silicon master pre-treated with silane (tridecafluro-1,1,2,2-tetrahydrooctyl-trichlorosilane, ABCR), then cured for 2 h at 65°C. After PDMS reticulation the inlets/outlet were drilled in the chip using a 0.75-mm diameter punch (Biopsy Punch, World Precision Instruments). The device was then assembled by covalently bonding the chip to a PDMS-coated glass coverslip (60 × 24 $mm^2$, bottom part) via oxygen-plasma treatment.

##### 1.3 Flow experiments and microscopy

*Microfluidic experiment.* The device inlets and outlet were connected via Teflon tubing (0.015" and 0.027" inner and outer diameters, Scientific Commodities) to a flow controller system (MFCS-8C, Fluigent) that allowed controlling the pressure drop between inlets and outlet and the circulation of cell suspensions in the microfluidic device. The in-slit pressure drop $\Delta P$ was calculated afterwards using COMSOL® software, from the inlet/outlet pressure drop and the number of open/clogged slits during the experiment Fig. S11. Devices with various series of slits (fixed width $W$, length $L$, and depth $D$ in each series) were placed on an inverted microscope (IX71, Olympus) equipped with cameras (high-speed camera Fastcam Mini, Photron and sCMOS camera Neo 5.5, Andor), 20x/60x/100x objectives, and temperature control. After BSA passivation of the device surfaces, RBCs were injected at 1% Hct at different temperatures and under different pressure drops in the physiological range (see Table S1).

*Observation of RBC retention and transit through slits.* RBCs were observed in brightfield with the 20× and 60× objectives for retention and transit quantification, respectively. Movies were acquired at high frame rate, from 500 to 1600 fps, with the Photron camera. In total, several tens of RBCs were imaged in all the experiments (see Table S1).

*Observation of RBC intracellular calcium.* RBCs were observed with the 100× objective. Movies were acquired in epi-fluorescence at 4 fps and exposure time of 250 msec with the Andor camera.

##### 1.4 Video analysis

Video analysis was mostly carried out manually using FIJI software. As a control before performing a complete analysis, the RBC projected radius was computed from an elliptic fit of the RBC projection. When RBCs reached the slits with abnormal shape (non discocyte) or radius (too large or too small), the data set was discarded.

*Retention analysis.* An RBC is considered retained if it stays in a slit more than 2 sec. When a slit is blocked by an RBC, those which arrive afterwards on the same slit are not counted. The retention rate is the ratio of the number of retained RBCs to the total number of RBCs (transiting and retained) which is at least 50. Retention is considered significant when it is higher than 10%.

*Transit analysis.* Using FIJI software the RBC position was tracked, and the RBC projected radius, shape and transit time were determined. RBCs were classified from their shape in two categories: "dumbbell" shape with rounded projections on each side of the slit; "tip" shape with a thin protrusion at the RBC front exiting from the slit. The manual processing was double-checked using Matlab routines. To measure automatically the RBC transit time, we designed an automated Matlab procedure to segment and track RBCs on live imaging. We first spatially registered images, from time-point to time-point, to isolate moving RBCs from the static background by subtracting the median projection of the overall movie from the movie itself. We then segmented RBCs using thresholding and applied a series of binary operations to improve the resulting object masks (holes filling, opening, erosion and size filtering). To reduce tracking mistakes, we also discarded aggregated RBCs (two or more touching RBCs) using a supervised KNN clustering algorithm considering features such as cell area and circularity. We finally followed individual RBCs over time by tracking overlapping binary objects from time point to time point. This allowed us to define the transit time as the time spent by a given RBC in contact with the slit.

*Intracellular calcium analysis.* The fluorescence intensity is measured over the RBC projected area and corrected by background subtraction. Calcium fluorescence is measured upstream of the slit when the RBC is not deformed, within the slit when the RBC is approximately symmetric in the slit, and downstream of the slit when the RBC has relaxed its shape.

##### 1.5 Statistics

Statistical tests are performed using the Mann-Whitney-Wilcoxon test in Kaleidagraph software (Synergy Software). Significant differences between data sets are indicated with stars: *, $p < 0.01$, **, $p < 0.001$, ***, $p < 0.0001$.

#### 2. Computational methods

##### 2.1 Determination of the standard RBC shape used in the simulations

Table S2 gives a survey of area and volume measurements in the literature. This table is made by appending latest data into a similar table made by Lim et al. (2). We used a standard shape with a surface area of 135 and the volume of 94, which is based on the original data by Evans and Fung (3) and very close to the latest data by Gifford et al. (4) and the average value of all existing data in Table S2. It is well known that there is some inaccuracy in the original data by Canham and Burton (5) since the edges of the RBCs cannot be clearly distinguished at that time and they are based on manual tracing of diametrical cross-sections of cells hanging vertically from the underside of microscope coverslips photographed edge-on. Comparing to other data, Canham and Burton (5) overestimated the volume. That is the main motivation of Evans and Fung (3) to improve the measurement of the geometry. Recently, Fischer (6) showed that the ratio of thickness across the dimple region to the thickness of the rim (THR) depends on the albumin concentration, and realized that the original work of Evans and Fung (3) underestimated this value, giving 0.314, significantly smaller than in the latest Fischer's work it was measured as 0.550 or 0.601 whether measured in plasma or serum. But for our current studies, as long the volume and surface are accuracy, the exact THR does not make any significant difference in terms of deformed cell shape or transit time, because the deformation is extremely large comparing to the initial shape. In addition, all the existing data are measured near room temperature, rather than body temperature. Based on the thermal expansion coefficient (1.2%/°C) of the red cell membrane measured by Waugh and Evans (7), we expect the surface area to be increased by 2% if the temperature is changed from 25°C to 37°C (12°C difference). Since the cell volume is determined by the osmotic pressure originating from the amount of hemoglobin molecules and their associated ions inside, the volume is not expected to change in isotonic solution since the hemoglobin amount is fixed. Since the critical pressure is highly sensitive to surface area, it is crucial to take the change of surface area due to temperature difference into account.

##### 2.2 Overview of the modeling approaches

While we applied an in-house code of coupled the finite element method (FEM) and boundary integral method to study the dynamics of RBCs passing through the slits with full fluid-structure interaction considered, we also applied the commercial FEM package ABAQUS (8) to study the retention of the RBCs passing through the slits as a quasi-static process without modeling the fluid flow explicitly. The reasons we used ABAQUS instead of the in-house code to study retention are: 1) Near the critical condition of retention, the process is so slow that it becomes too long for the in-house code to simulate, while it approaches a quasi-state process so that ABAQUS can simulate the process much more efficiently and accurately; 2) For the retention study, we are only interested in the critical pressure for the cells to pass rather than its dynamics, so that it is not necessary to apply the in-house code for dynamics; 3) To accurately predict the critical pressure, the experimental measured area modulus of the lipid bilayer (375-450 mN/m) (7) must be used, because the area deformation plays a significant role on the critical pressure for retention which must be accurately captured even it is small. This large bilayer area modulus significantly decreases the stable time step size of the simulation, which makes the fluid-structure simulations using the in-house code prohibitively expensive. For typical simulations of the dynamics of RBCs passing through the slits, since it is far away from the critical retention condition, we used lower bilayer area modulus in the in-house code for the fluid-structure simulations to allow larger time step size but still limit the local area deformation within 1%.

##### 2.3 ABAQUS simulations of RBC retentions

We used ABAQUS explicit solver (8) to simulate the passing of RBCs through the slit as a quasi-static process by adding artificial mass and damping. We applied 8878 S4 shell elements to simulate the RBC membrane. We did systematic convergence test to make sure the final results (such as critical pressure) do not depend on mesh size and artificial mass and damping. We implemented the multiscale model (see §2.4 below) of the RBC membrane in ABAQUS using the user subroutine VUMAT. The slit walls were modeled as rigid surfaces based on experimentally measured dimensions. The pressure was applied on the front half surface of the cell to push it through the slit. General contacts were imposed between all the surfaces. We adopted fully 3D simulations rather than utilizing the symmetry, because buckling of the cell membrane can occur which breaks the symmetry. Surface-based fluid cavity was used to enforce the volume conservation by assigning an apparent fluid bulk modulus. The apparent bulk modulus of the fluid due to osmotic pressure was estimated by following the approach used in Evans and Waugh (9):
The ratio of the volume change $\Delta V$ to the initial volume $V_0$ can be related to the pressure difference as

$$\frac{\Delta V}{V_0} = -\frac{R_w \Delta P}{\beta C_i}, \qquad [1]$$

where $\beta = 310\text{K} \cdot 8.31\text{J/K/mol} = 2.5775 \times 10^3$ J/mol is the gas constant times absolute temperature, $C_i = 300$ mM $= 300 \times 10^{-3}$ mol/L $= 300$ mol/m$^3$ is the molar concentration of solute species inside the cell, including 150 mM from $K^+$ and 150 mM from amino acids, and $R_w = 0.6$ is the fraction of total cell volume that is water. Thus, the apparent fluid bulk modulus $B_f = \frac{\beta C_i}{R_w} = 1.289 \times 10^6$ Pa.

##### 2.4 Multiscale modeling of RBC membranes and spectrin unfolding.

A multiscale RBC model was implemented in both the in-house code and an ABAQUS VUMAT user subroutine for modeling material properties (8). In the whole-cell continuum-level model, the Cauchy stress resultant is given as

$$\boldsymbol{\sigma}^{\boldsymbol{s}}(\mathbf{F}) \cdot h = \tau_m(\alpha,\beta)\boldsymbol{I} + \frac{\mu(\alpha,\beta)}{(\alpha+1)^2}(\boldsymbol{B} - \frac{trace(\boldsymbol{B})}{2}\boldsymbol{I}), \qquad [2]$$

where $\tau_m$ and $\mu$ are the membrane mean stress resultant and shear modulus. $\boldsymbol{B} = \boldsymbol{F}\boldsymbol{F}^T$ is the left Cauchy-Green deformation tensor and $h$ is the thickness of the membrane modeled as a thin shell. $\boldsymbol{F}$ is the deformation gradient. $I$ is the identity matrix, $\alpha = \lambda_1\lambda_2 - 1$ and $\beta = (\lambda_1^2 + \lambda_2^2)/(2\lambda_1\lambda_2) - 1$ are two invariants representing the area and shear deformation respectively (7). $\lambda_1$ and $\lambda_2$ are the principal stretches, which can be calculated from $\boldsymbol{F}$.

Details of the formulation of the multiscale model, including the expressions of $\tau_m$ and $\mu$, can be found in Ref. (10), in which the distributions of the spectrin orientation and natural length from cyro-EM data (11) and the updated copy numbers of spectrin tetramers from proteomic data (12, 13) are taken into account. One major difference here is that we included spectrin unfolding in the current study, while it was ignored in Ref. (10). We considered a spectrin tetramer with $n_f$ folded domains with a domain length of $r_f$ and $n_u$ unfolded domains with a domain length of $r_u$, the total length $s$ of a spectrin tetramer is given as (14, 15)

$$s = n_f r_f + n_u r_u. \qquad [3]$$

Dividing the equation by the total number of spectrin domains $n = n_f + n_u$ in a tetramer and the undeformed domain length of $r_0$, we have the spectrin deformation $x = \frac{s}{nr_0} = (1-\phi_u)\frac{r_f}{r_0} + \phi_u\frac{r_u}{r_0}$, where the fraction of unfolded domains $\phi_u = n_u/n$ can be calculated as

$$\phi_u = \frac{\exp\frac{(F-F_{1/2})\Delta\Delta x^*}{k_B T}}{1-\exp\frac{(F-F_{1/2})\Delta\Delta x^*}{k_B T}}, \qquad [4]$$

where $\Delta\Delta x^*$ is the combined activation length of the unfolding and refolding processes, $F_{1/2}$ is the force by which half of the spectrin domains are unfolded (14, 15), $k_B$ is the Boltzmann constant and $T$ is the absolute temperature. $r_f$ and $r_u$ are related to the force $F$ using worm-like chain model (16) with different persistence lengths ($p_f$ and $p_u$) and contour lengths

($L_f$ and $L_u$), respectively. For specific values, we used $\Delta\Delta x^* = 12.6$ nm, $L_f = 4$ nm, $L_u = 39$ nm, $p_f = 20$ nm, $p_u = 0.8$ nm, $F_{1/2} = 1$ pN at 37°C and $F_{1/2} = 10$ pN at 25 °C. The spectrin orientation distribution and the natural length distribution used in this study are the same as our previous study in Ref. (10), which gives an initial shear modulus $\mu_0$ about 9 pN/$\mu$m. The membrane viscoelasticity is modeled as a Voigt model with a characteristic time $t_c = \eta_m/\mu_0$ as 0.1 s as measured in experiments (17), where $\eta_m = 0.9$ pN·s/µm is the membrane viscosity and $\mu_0$ is the initial value of $\mu$ without deformation. The detailed numerical implementation of this viscoelastic model can be found in (18).

#### 3. Surface area and volume of an RBC in a slit approximated as two tether-connected cone-spheres

The microfluidic device has slanted walls at the entrance/exit of the slits making an angle of 36.25° with the direction of the flow, which does not allow a spherical deformation in the immediate vicinity of the slit. This is a more demanding geometry for the $S/V$ ratio. For this geometry, there is no simple analytical solution allowing to calculate the minimal surface of an RBC with a given volume to pass in the considered slit. Therefore, as a first approximation, we considered a cone-sphere geometry and calculated the surface area as a function of the volume from a conical shape completed by a spherical cap. In the middle of a slit, the RBC is described by two symmetrical cone-spheres connected by an infinitely thin tether.

The volume $V_C$ and surface $S_C$ of a cone are:

$$V_C = \frac{\pi}{3} R^3 \frac{cos\theta^4}{sin\theta} \tag{5}$$

and

$$S_C = \pi R L = \pi R^2 \frac{cos\theta^2}{sin\theta}, \tag{6}$$

where $R$ is the radius of the cone base, $\theta$ is the angle between the walls at the edge of the slit and the direction of flow, and $L$ is the length of the slit.

The volume $V_{Sph}$ and surface $S_{Sph}$ of a spherical cap of radius $R$ are:

$$V_{Sph} = \frac{\pi R^3}{3}(1 + sin\theta)^2(2 - sin\theta)^2 \tag{7}$$

and

$$S_{Sph} = 2\pi R D = 2\pi R^2(1 + sin\theta). \tag{8}$$

The volume $V_{CS}$ and surface $S_{CS}$ of a cone-sphere are:

$$V_{CS} = \frac{\pi R^3}{3}(1 + sin\theta)^2(2 - sin\theta) + \frac{\pi R^3}{3}\frac{cos\theta^4}{sin\theta} \tag{9}$$

and

$$S_{CS} = \pi R^2(2(1 + sin\theta) + \frac{cos\theta^2}{sin\theta}). \tag{10}$$

With $\theta = 36.25°$, $V_{CS}$ and $S_{CS}$ express as:

$$V_{CS} = 4.31\frac{\pi R^3}{3} \tag{11}$$

and

$$S_{CS} = 4.31\pi R^2. \tag{12}$$

The total area $S$ and volume $V$ of the two cone-spheres are thus:

$$S = 2S_{CS} = 8.62\pi R^2 \tag{13}$$

and

$$V = 2V_{CS} = 8.62\frac{\pi R^3}{3}. \tag{14}$$

The relation between $S$ and $V$ is therefore:

$$S = (8.62\pi \times 9)^{1/3} \times V^{2/3} = 6.25 \times V^{2/3}. \tag{15}$$

| Slit dimensions, µm$^3$ | Temperature, °C | In-slit pressure drop, Pa | Retention rate, % (N) | Transit time, ms (N') |
|---|---|---|---|---|
| 0.43×2.10×4.90 | 21 | 191 | 16 (111) | - |
| 0.42×2.20×5.40 | 21 | 394 | 23 (133) | - |
| 1.09×2.59×4.80 | 15 | 489 | - | 80.0 (41) |
| | | 997 | - | 48.0 (4) |
| | | 1375 | - | 29.0 (15) |
| | 37 | 150 | - | 10.0 (29) |
| | | 187 | - | 6.30 (73) |
| | | 247 | - | 5.60 (44) |
| | | 560 | - | 2.40 (28) |
| | | 660 | - | 1.90 (30) |
| 0.94×2.28×5.30 | 15 | 224 | - | 28.0 (68) |
| | | 448 | - | 12.0 (80) |
| | | 1256 | - | 4.0 (57) |
| 0.94×2.40×5.20 | 22 | 231 | - | 17.0 (24) |
| | | 462 | - | 6.90 (60) |
| 0.90×2.20×5.30 | 15 | 485 | - | 13.0 (55) |
| | | 967 | - | 7.40 (25) |
| | | 1459 | - | 6.40 (19) |
| 0.87×2.26×4.80 | 37 | 192 | - | 12.0 (53) |
| | | 331 | - | 6.90 (77) |
| | | 500 | - | 4.70 (26) |
| 0.86×2.75×4.70 | 15 | 360 | - | 33.0 (80) |
| | | 375 | - | 24.0 (59) |
| | | 735 | - | 12.0 (60) |
| | 22 | 445 | - | 9.40 (35) |
| | | 885 | - | 3.80 (62) |
| | | 1500 | - | 2.20 (28) |
| | 24 | 216 | - | 32.0 (32) |
| | | 445 | - | 8.10 (39) |
| | | 865 | - | 3.80 (18) |
| | 37 | 218 | - | 17.0 (23) |
| | | 440 | - | 6.30 (25) |
| | | 835 | - | 3.10 (11) |
| | | 280 | - | 11.0 (35) |
| | | 695 | - | 3.70 (44) |
| | | 1200 | - | 1.90 (46) |
| 0.68×2.70×4.70 | 37 | 450 | - | 11.0 (19) |
| | | 894 | - | 5.00 (21) |
| | | 1700 | - | 2.40 (24) |
| 0.67×1.90×4.70 | 22 | 190 | - | 48.0 (15) |
| | | 360 | - | 24.0 (10) |
| | | 775 | - | 9.40 (61) |
| 0.61×2.01×5.0 | 37 | 325 | - | 22.2 (44) |
| | | 645 | - | 9.0 (29) |
| | | 1325 | - | 5.0 (60) |
| 0.61×2.01×5.0 (1mM diamide) | 37 | 325 | - | 24.5 (65) |
| | | 645 | - | 12.0 (71) |
| | | 1325 | - | 6.50 (81) |
| 0.61×2.01×5.0 (400 mOsm) | 37 | 325 | - | 28.0 (65) |
| | | 645 | - | 13.0 (47) |
| | | 1325 | - | 8.2 (37) |
| 0.61×2.01×5.0 (500 mOsm) | 37 | 325 | - | 43.4 (47) |
| | | 645 | - | 23.2 (54) |
| | | 1325 | - | 10.3 (47) |
| 0.45×2.21×5.40 | 37 | 63 | - | 1830.0 (3) |
| | | 126 | - | 350.0 (44) |
| | | 190 | - | 160.0 (69) |
| | | 317 | - | 71.0 (89) |
| | | 633 | - | 37.0 (51) |
| | | 220 | - | 120.0 (21) |

| | | | | |
|---|---|---|---|---|
| | | 366 | - | 100.0 (33) |
| | | 728 | - | 37.0 (180) |
| | | 1456 | - | 15.0 (215) |
| 0.37×1.40×4.70 | 37 | 195 | - | 250.0 (43) |
| | | 312 | - | 110.0 (62) |
| | | 408 | - | 69.0 (114) |
| 0.30×1.88×4.80 | 37 | 490 | - | 160.0 (87) |
| | | 820 | - | 81.0 (42) |
| | | 1630 | - | 38.0 (110) |
| 0.28×1.87×5.00 | 37 | 490 | - | 461.0 (54) |
| | | 815 | - | 217.0 (10) |
| | | 1623 | - | 84.0 (10) |

**Table S1. Summary of performed experiments:** Experimental conditions (slit dimensions, temperature, in-slit pressure drop) and quantification (retention rate when RBCs are blocked in the slits, median transit time when RBCs pass through slits). N,N': number of analyzed RBCs.

| | Surface area ($\mu m^2$) | Volume ($\mu m^3$) | 2% extra area (37 °C) | Sphericity (25 °C) | Sphericity (37 °C) |
|---|---|---|---|---|---|
| Canham and Burton (1968) (5) | 138.1 | 107.5 | 140.862 | 0.791707799 | 0.776184117 |
| Evans and Fung (1972) (3) | 135 | 94 | 137.7 | 0.740578488 | 0.726057341 |
| Jay (1975) (19) | 136.9 | 104.2 | 139.638 | 0.782218332 | 0.766880718 |
| Jay (1976) (19) | 133.4 | 98.1 | 136.068 | 0.771098398 | 0.755978822 |
| Fung et al. (1981) (20) | 129.95 | 97.91 | 132.549 | 0.790547632 | 0.775046698 |
| Linderkamp and Meiselman (1982) (21) | 134.1 | 89.8 | 136.782 | 0.723172227 | 0.708992379 |
| Nash and Meiselman (1983) (22) | 137 | 99 | 139.74 | 0.755421234 | 0.740609053 |
| Linderkamp et al. (1983) (23) | 134.4 | 88.4 | 137.088 | 0.714038891 | 0.700038128 |
| Linderkamp et al. (1986) (24) | 132.1 | 94.9 | 134.742 | 0.761659651 | 0.746725148 |
| Stadler and Linderkam (1989) (25) | 137.1 | 90.5 | 139.842 | 0.711019028 | 0.697077479 |
| Waugh et al. (1992) (26) | 135 | 93 | 137.7 | 0.735316801 | 0.720898825 |
| Linderkamp et al. (1993) (27) | 137.1 | 90.5 | 139.842 | 0.711019028 | 0.697077479 |
| Engström and Löfvenberg (1998) (28) | 141.4 | 105.7 | 144.228 | 0.764575201 | 0.74958353 |
| Ruef and Linderkamp (1999) (29) | 132 | 101 | 134.64 | 0.79455984 | 0.778980235 |
| Gifford et al. (2003) (4) | 134 | 95 | 136.68 | 0.751387381 | 0.736654295 |
| **Average** | **135.17** | **96.634** | **137.8734** | **0.753400579** | **0.738628019** |

**Table S2. Area and volume measurements in the literature.**

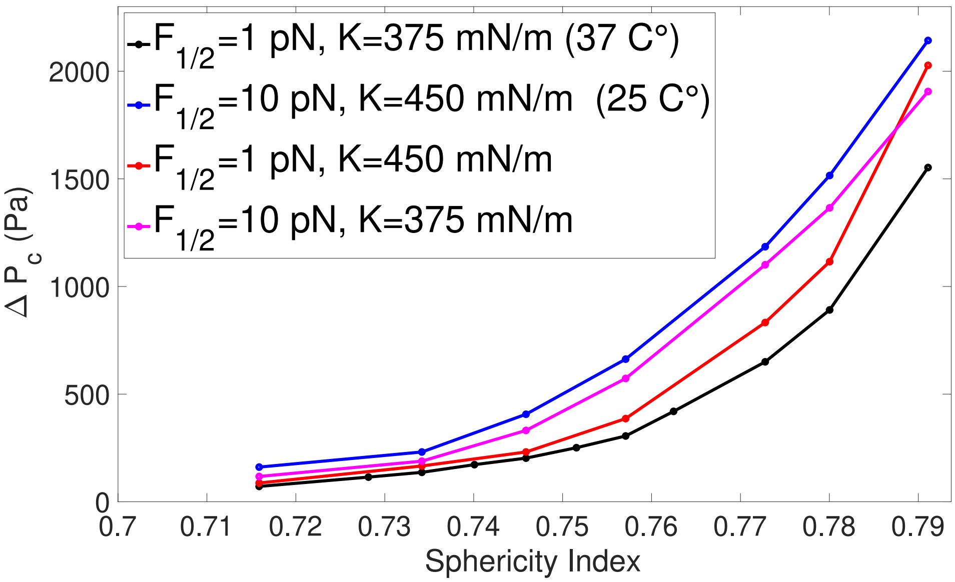


**Fig. S1.** Effects of temperature on the relation between the sphericity index and the critical pressure required for an RBC to pass through the reference slit (0.37×1.30×4.70 µm$^3$). $F_{1/2}$ is the force to unfold half of the spectrin domains, and K is the bilayer area modulus.

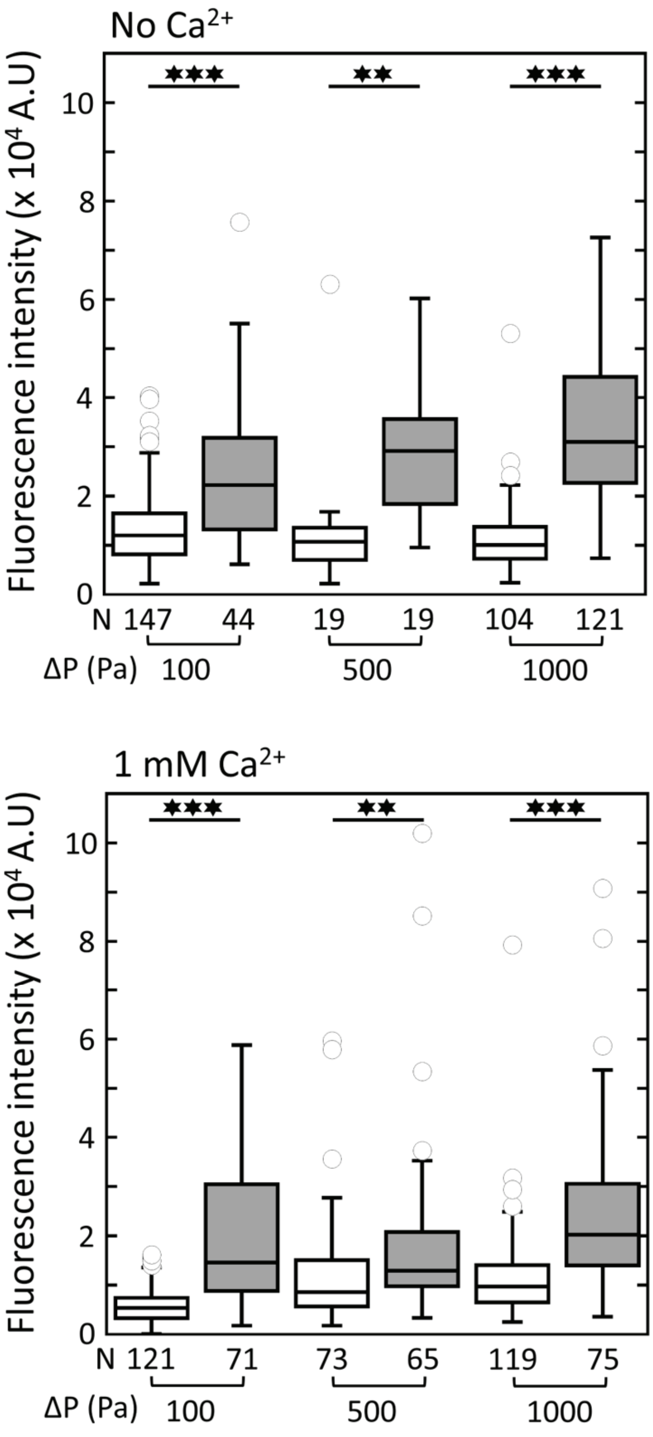


**Fig. S2.** Intracellular calcium measured in RBCs upstream (white) and downstream (grey) of 0.80×2.78×4.70 µm$^3$ slits under in-slit pressure drops $\Delta P$ of 100, 500, and 1000 Pa in the absence (top) and presence (bottom) of 1 mM calcium in the RBC suspension buffer. N: number of analyzed RBCs. **: p < 0.001, ***: p < 0.0001.

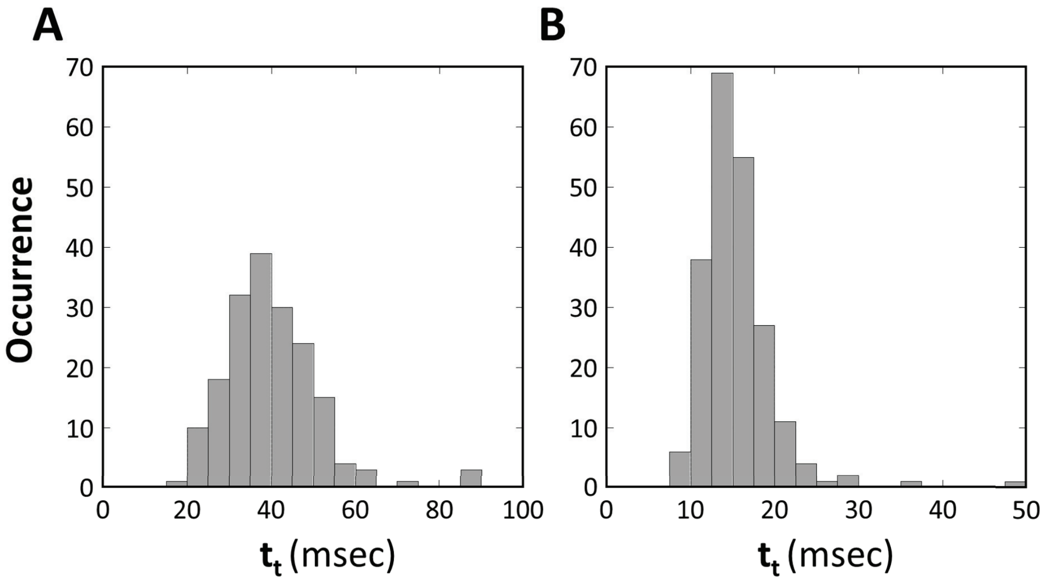


**Fig. S3.** Distribution of RBC transit times $t_t$ measured in a 0.45×2.21×5.4 µm$^3$ slit under in-slit pressure drops of (A) 728 Pa and (B) 1456 Pa at 37°C. Number of analysed RBCs: (A) 180 and (B) 215.

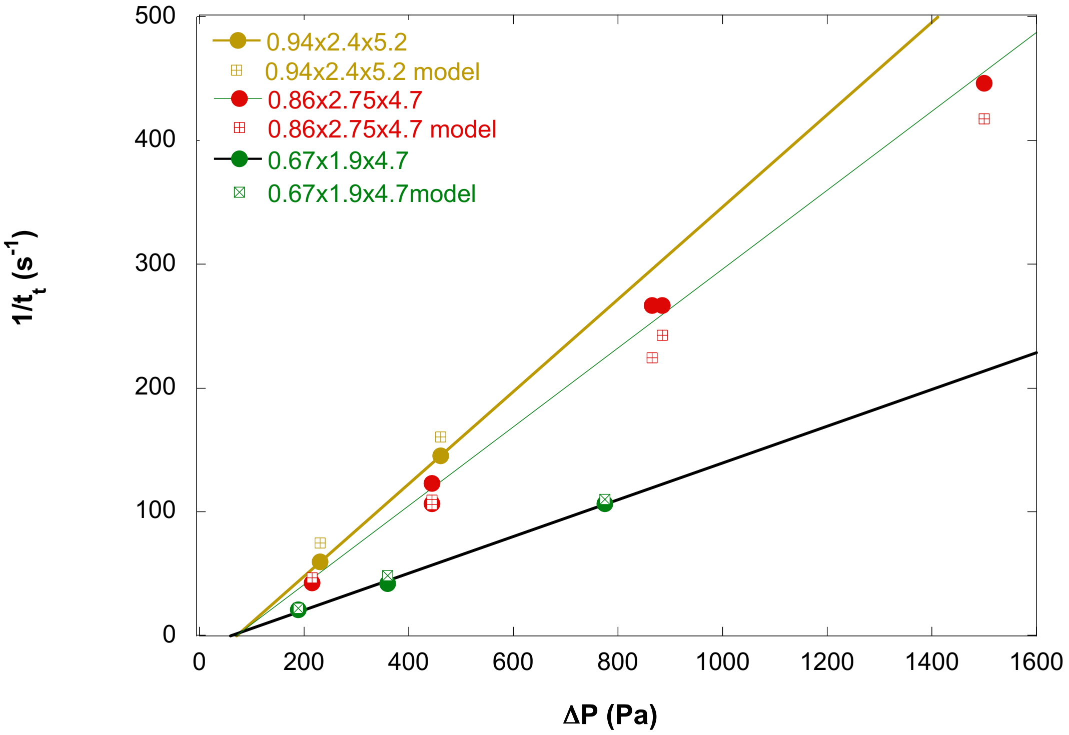


**Fig. S4.** Inverse of the transit time $1/t_t$ versus in-slit pressure drop $\triangle$P, experiments and simulations at 22℃.

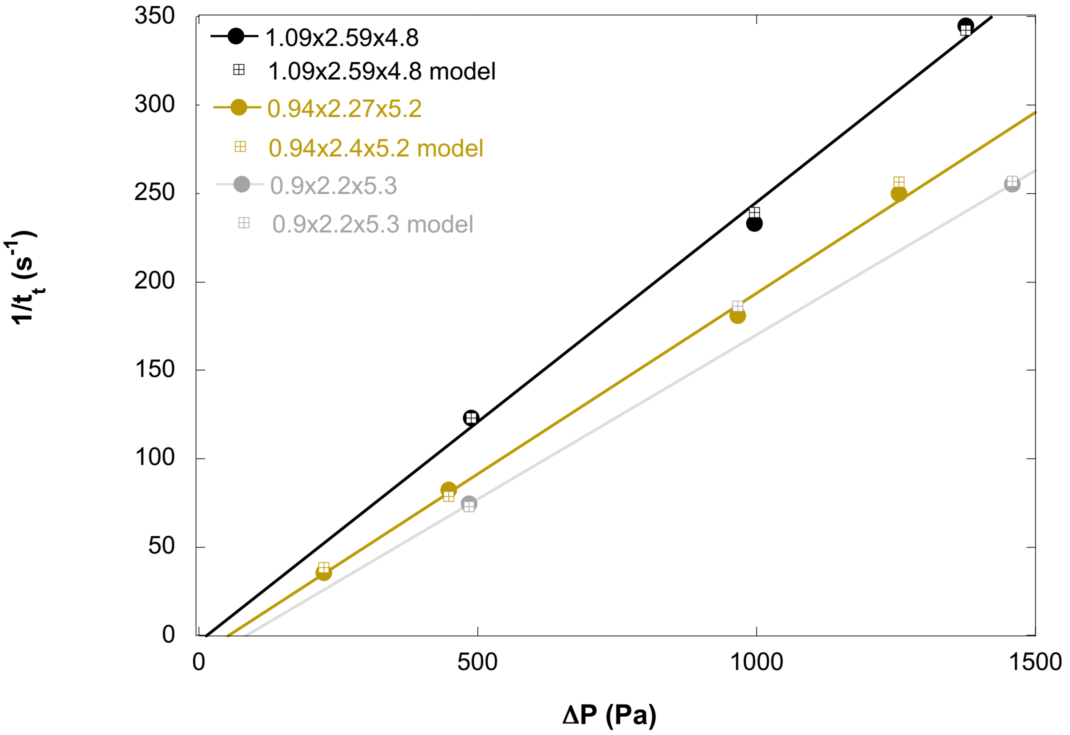


**Fig. S5.** Inverse of the transit time $1/t_t$ versus in-slit pressure drop $\triangle$P, experiments and simulations at 15℃.

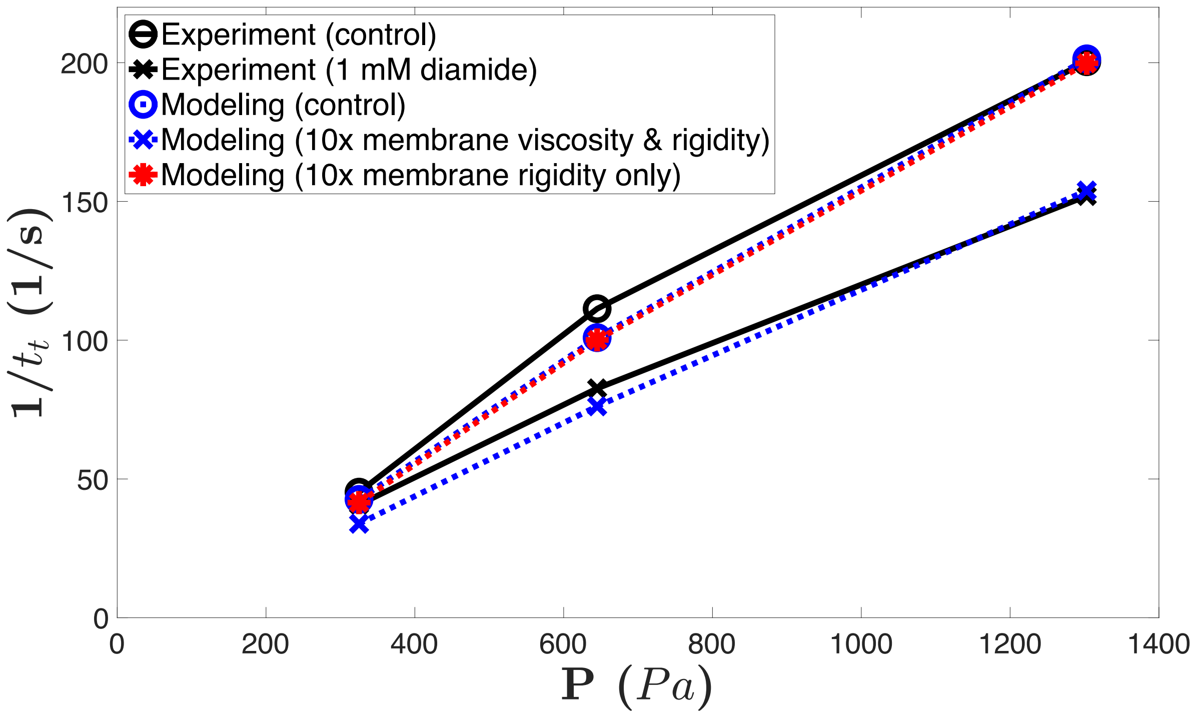


**Fig. S6.** Inverse of the transit time $1/t_t$ versus in-slit pressure drop ΔP in $0.61 \times 2.01 \times 5.0$ µm$^3$ slits at isotonic osmolarity without and with 1mM diamide. Numerical simulations fit well the experimental data. For the control case, the membrane viscoelasticity is modeled using a multiscale model described in the *SI Appendix*, section 2.4, giving an initial cytoskeleton shear modulus $\mu_0$ = 9 pN/µm. The characteristic time $t_c = \eta_m/\mu_0$ is chosen as 0.1 s as measured in experiments (17), where $\eta_m$ = 0.9 pN·s/µm is the membrane viscosity. For the diamide-treated case, we first increased both the initial shear modulus and membrane viscosity by 10 times (30), while keeping $t_c$, $F_{1/2}$, and $K$ unchanged. This leads to slower passage (blue curve with crosses). We then increased only the membrane shear modulus by 10 times, and it didn't slow down the passage, and the transit time remains almost the same as the control case (red curve). Lines are guide to the eye.

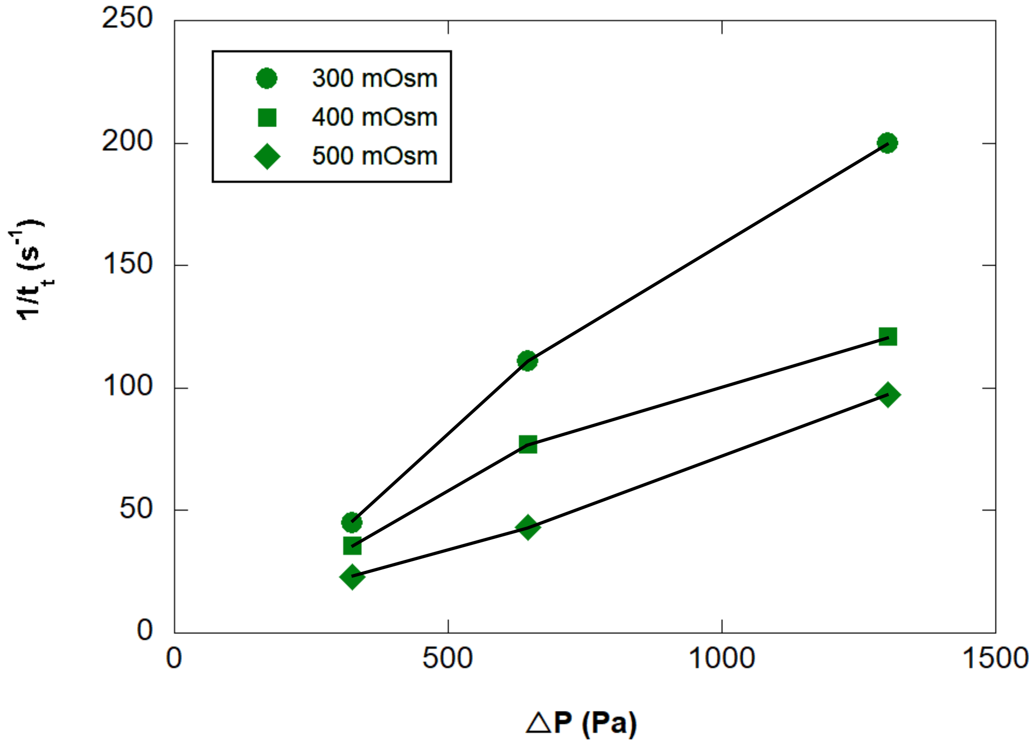


**Fig. S7.** Inverse of the transit time $1/t_t$ versus in-slit pressure drop ΔP in $0.61 \times 2.01 \times 5.0$ µm$^3$ slits upon increasing osmolarity of RBC suspension buffer at 37 °C. Isotonic condition is 300 mOsm. Lines are guide for the eye.

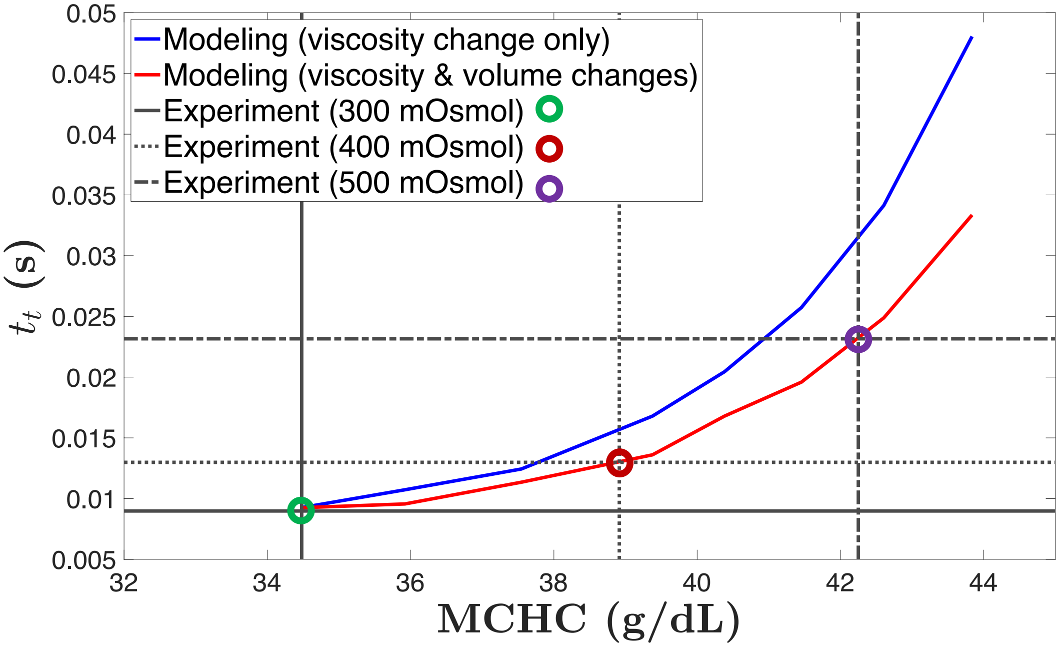


**Fig. S8.** Transit time $t_t$ as functions of altered mean corpuscular hemoglobin concentration (MCHC) due to varying osmolarity. The blue curve is the computational prediction of the transit time with only viscosity change based on altered MCHC using the Ross and Minton model (31) based on the experimental measurements (32–34). For the red curve, besides viscosity change, the cell volume reduction is also considered. Volume reduction is calculated based on constant hemoglobin amount inside a cell. The horizontal lines are experimental measured transit time under three different osmolarities. The vertical lines are the corresponding MCHC predicted by computational modeling for the experimentally measured transit time. The slit dimensions are $0.61 \times 2.01 \times 5.0$ µm$^3$ and the in-slit pressure is 645 Pa.

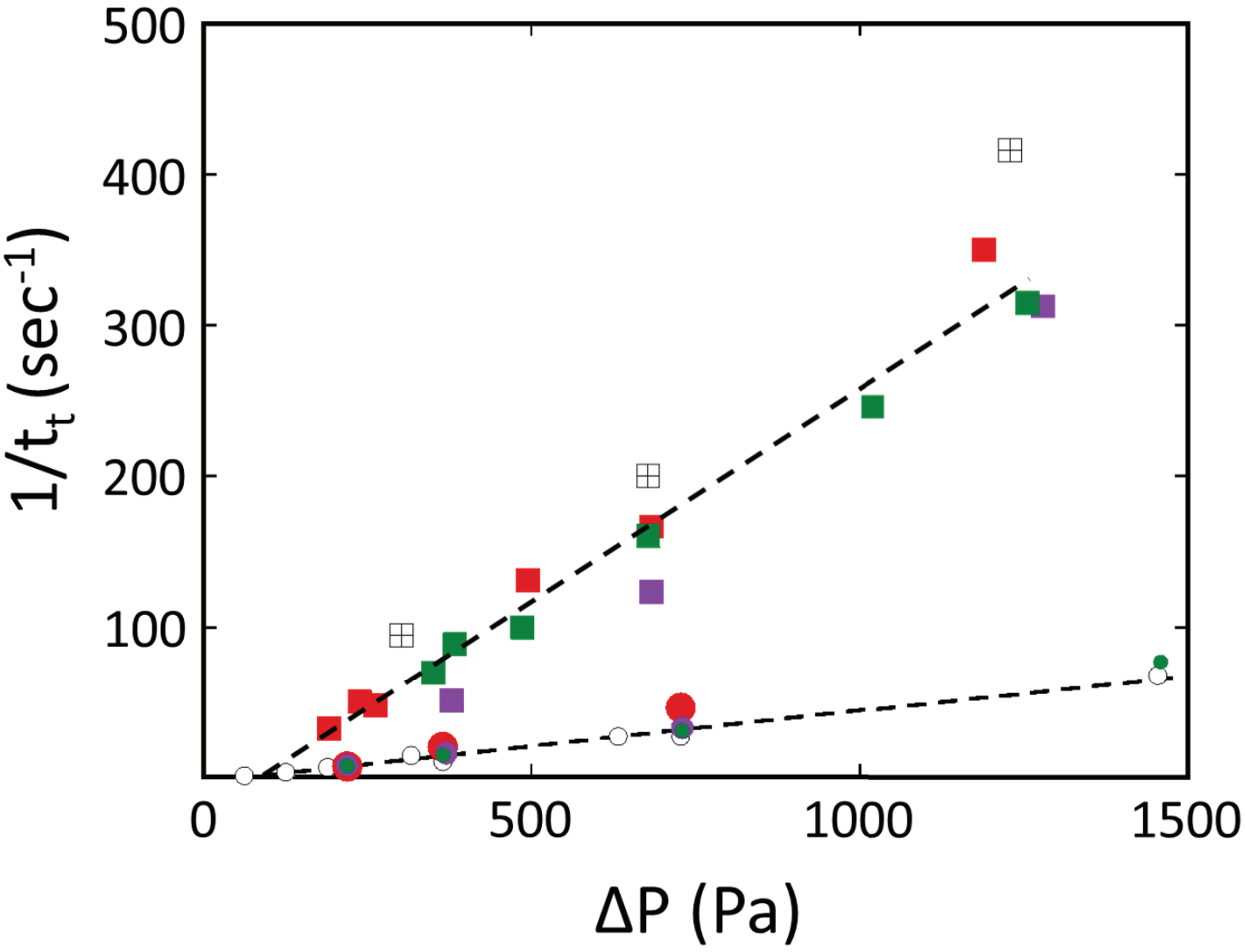


**Fig. S9.** Inverse of the transit time $1/t_t$ versus in-slit pressure drop $\triangle$P in 0.80-µm (square symbols) and 0.45-µm (circle symbols) wide slits in the absence or presence of 1 mM calcium in the suspension buffer (open and closed symbols, respectively). The black open squares with cross correspond to no-calcium condition in presence of 10% plasma in the suspension buffer. In calcium conditions, the suspension buffer always contains 10% plasma, and is supplemented with inhibitors of PIEZO1 (5 µM GsMTX4, in purple) or Gardos channels (10 µM TRAM34, in green) or not (in red). In all conditions, $1/t_t$ varies linearly with $\triangle$P.

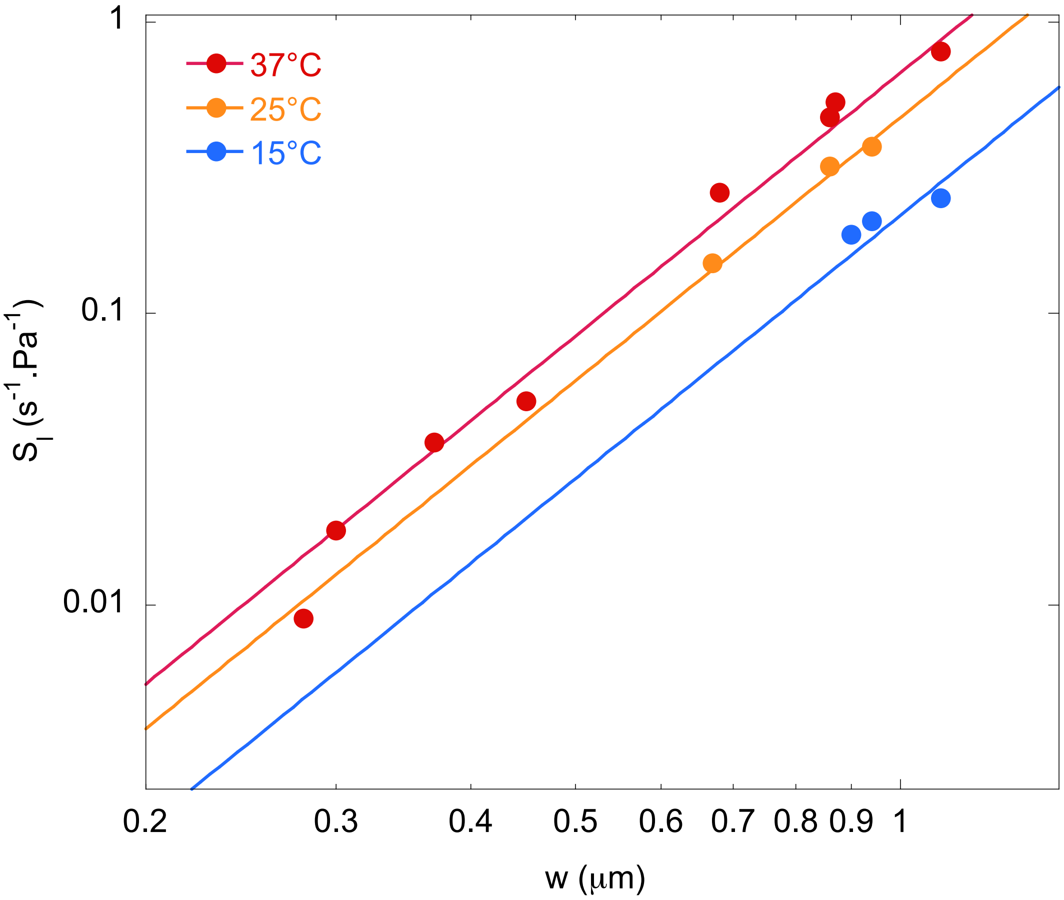


**Fig. S10.** Log-log representation of the slope of the $1/t_t$ variation vs $\Delta$P with the slit width $W$ at 37℃, 22℃, and 15℃. Solid lines are fits by a power law with the exponent 3.

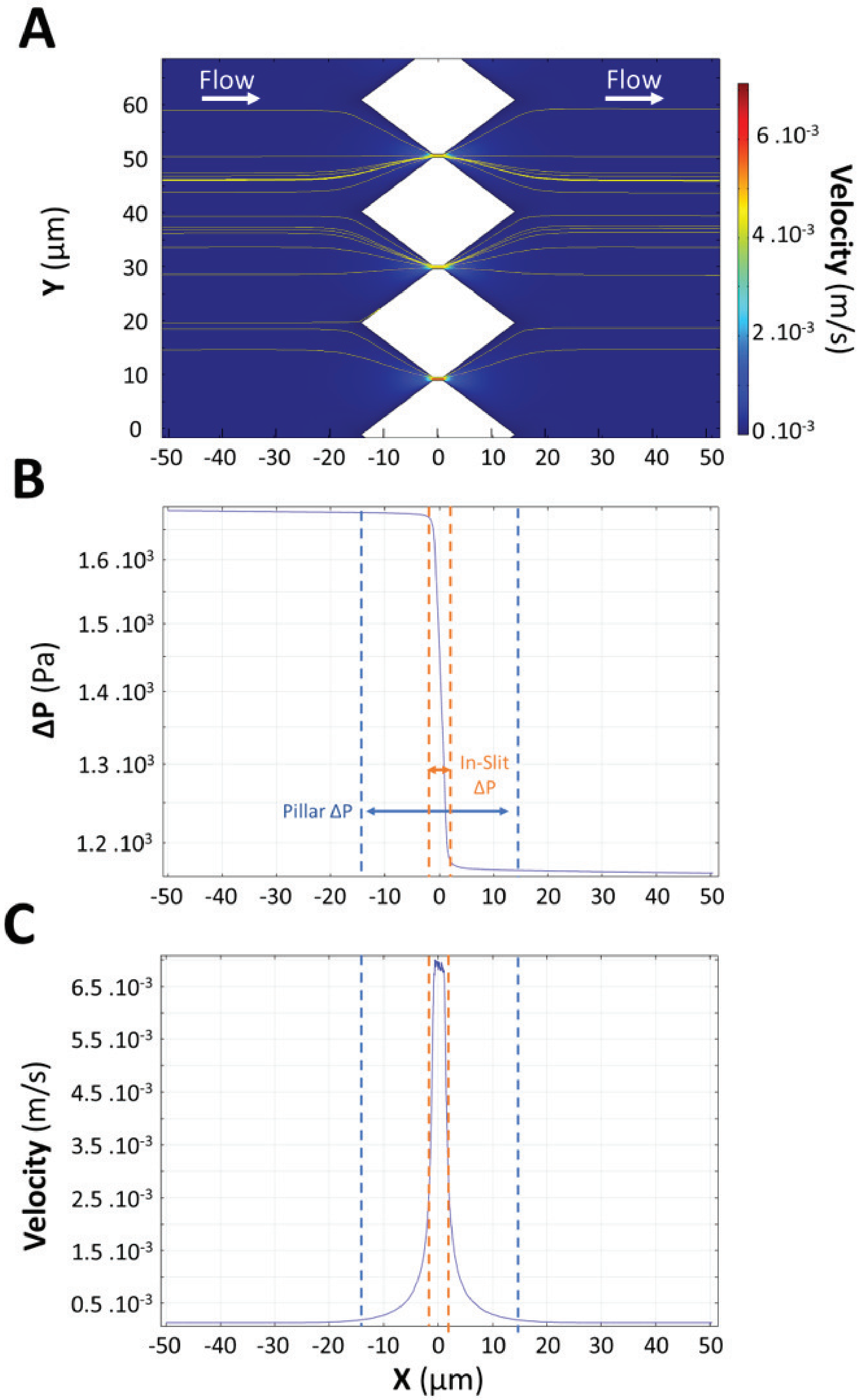


**Fig. S11.** ComsolR© calculation of streamlines, pressure drop, and fluid velocity over a region spanning over 50 µm from each side of the slits. (A) Image showing the streamlines in the slits. The color code indicates the velocity magnitude. (B-C) Evolution of pressure drop ΔP (B) and fluid velocity (C) as a function of the position X in the channel. The dashed lines indicate the borders of the pillars (blue) and the slit (orange). The position X = 0 µm corresponds to the center of the slits.

**Movie S1. Movie of an RBC passing through a $0.28 \times 1.87 \times 5.00$ µm$^3$ slit.** Time is in msec. Scale bar: 10 µm.

**Movie S2. Movie of an RBC displaying a dumbbell shape while passing through a $0.94 \times 2.28 \times 5.30$ µm$^3$ slit.** Time is in msec. Scale bar: 10 µm.

**Movie S3. Movie of an RBC displaying a front tip while passing through a $0.94 \times 2.28 \times 5.30$ µm$^3$ slit.** Time is in msec. Scale bar: 10 µm.

**Movie S4. Movie of the 3D-view of an RBC in a $0.67 \times 2.34 \times 4.98$ µm$^3$ slit.** The movie starts from a top view of the RBC in the slit whose length and width only are visible. The rotation of the RBC shows the thin neck between the RBC parts on each side of the slit and the slanted shape of the RBC close to the slit due to the oblique slit walls. Scale bar: 3 µm. The RBC membrane is labelled with CellTrace Yellow.

**Movie S5. Movie of a spherocytic RBC blocked in a $0.63 \times 2.70 \times 5.0$ µm$^3$ slit.** Time is in sec. Scale bar: 10 µm.

**Movie S6. Movie of an irreversible sickle RBC passing through a $0.63 \times 2.70 \times 5.0$ µm$^3$ slit.** Time is in sec. Scale bar: 10 µm.
